# An INDEL genomic approach to explore population diversity of phytoplankton

**DOI:** 10.1101/2023.02.09.527951

**Authors:** Martine Devic, Louis Dennu, Jean-Claude Lozano, Cédric Mariac, Valérie Vergé, Philippe Schatt, François-Yves Bouget, François Sabot

## Abstract

**Background:** Although metabarcoding and metagenomic approaches have generated large datasets on worldwide phytoplankton species diversity, the intraspecific genetic diversity underlying the genetic adaptation of marine phytoplankton to specific environmental niches remains largely unexplored. This is mainly due to the lack of biological resources and tools for monitoring the dynamics of this diversity in space and time.

**Results:** To gain insight into population diversity, a novel method based on INDEL markers was developed on *Bathycoccus prasinos* (Mamiellophyceae), an abundant and cosmopolitan species with strong seasonal patterns. Long read sequencing was first used to characterise structural variants among the genomes of six *B. prasinos* strains sampled from geographically distinct regions in the world ocean. Markers derived from identified insertions/deletions were validated by PCR then used to genotype 55 *B. prasinos* strains isolated during the winter bloom 2018-2019 in the bay of Banyuls-sur-Mer (Mediterranean Sea, France). This led to their classification into eight multi-loci genotypes and the sequencing of strains representative of local diversity, further improving the available genetic diversity of *B. prasinos*. Finally, selected markers were directly tracked on environmental DNA sampled during 3 successive blooms from 2018 to 2021, showcasing a fast and cost-effective approach to follow local population dynamics.

**Conclusions:** This method, which involves (i) pre-identifying the genetic diversity of *B. prasinos* in environmental samples by PCR, (ii) isolating cells from selected environmental samples and (iii) identifying genotypes representative of *B. prasinos* diversity for sequencing, can be used to comprehensively describe the diversity and population dynamics not only in *B. prasinos* but also potentially in other generalist phytoplankton species.

## Background

Marine phytoplankton, including picoeukaryotic algae, is responsible for a large fraction of primary production [1]. In temperate regions, the abundance and diversity of the phytoplankton is often seasonal and occurs in bursts, as algal blooms. *Per se*, blooms have a large impact on global primary production and therefore the understanding of the genetic basis of phytoplankton adaptation to seasonal niches and the effects of ocean warming on phytoplankton blooms are of the utmost importance.

Meta-ribosomal barcoding on the nuclear or plastidial 18/16S rRNA gene has opened the access to massive data in time and space and has accelerated the study of phytoplankton species and genera diversity in natural communities. However, metabarcoding approaches do not provide information on intraspecific genetic variations. Assessing only interspecific diversity leads to the underestimation of the diversity of the populations. Equally, a single isolate cannot represent the diversity of a population. Since natural selection acts on variation among individuals within populations, it is essential to incorporate both intra and interspecific trait variability into community ecology [2]. Indeed, Raffard et al. [3] demonstrated that intraspecific variation has significant ecological effects across a large set of species, confirming a previous estimate based on a more restricted species set [4]. Furthermore, it has been shown that diversity within species is rapidly decreasing, making them more homogeneous and highlighting the need to preserve intraspecific variations since this intraspecific diversity reinforces the overall population stability in the face of environmental change [4].

In diatoms, intraspecific variations play a key role in the response of species to several important environmental factors such as light, salinity, temperature and nutrients [5]. Modelling approaches suggest that intraspecific variation extends bloom periods by providing variability in competitive interactions between species under changing conditions, and contributes to fitness in temporary microhabitats. Intraspecific genetic diversity allows local adaptation by optimising physiological responses, allowing local populations to be exceptionally fit and competitive in their respective habitat.

Since an assembly of genotypically diverse individuals constitutes a population, methodological approaches have been developed in order to determine the genetic variation among individuals [6]. One of the main challenges is the difficulty of isolating a sufficient number of individuals for classical diversity analyses while ensuring representativity of diverse subpopulations. Historically, sequence-based intraspecific markers correspond to small nucleotide repeats such as microsatellites [7], chloroplastic [8] and mitochondrial [9] genes and a few nuclear genes that were applied to several hundreds of isolated individuals. The development of Randomly Amplified Polymorphic DNA (RAPD) markers [10] and more recently of Restriction-site-Associated DNA sequencing techniques (RADseq) [11] allowed the discovery and genotyping of thousands of genetic markers for any given species at relatively low-cost. Some of these approaches have been used to analyse the diversity of populations during algal blooms [12]. Recently, the dramatic increase of the number of sequenced genomes led to large-scale diversity studies with large sets of nuclear genes or whole genome comparisons. However, most of these approaches require the isolation of a large number of individuals. As a consequence, the intraspecific diversity remains poorly documented in marine phytoplankton.

Widely distributed from the equator to arctic and antarctic poles with a marked seasonality in temperate and polar regions, picoeukaryotes belonging to the *Mamiellales* order (*Bathycoccus*, *Ostreococcus* and *Micromonas*) are abundant and have a cosmopolitan presence, illustrating a high capacity for adaptation to a wide range of contrasting environments [13–16]. Novel, rapid and cheap sequencing technologies have given access to *Mamiellales* interspecific diversity by metagenomic [16,17] and metatranscriptomic approaches [18]. However, to date, very little information is available on intraspecific diversity of *Bathycoccaceae*, with the exception of *Ostreococcus tauri* from mediterranean lagoons [19]. Unlike *O. tauri*, which is usually not detectable in publicly available metagenomes, *Bathycoccus sp.* is the most cosmopolitan *Mamiellophyceae* genus. Although *Bathycoccus sp.* is abundant in the world Ocean, Metabarcoding of 18S ribosomal RNA (V9 region) does not allow the identification of *Bathycoccus* species [20,21]. Through metagenomic analysis, *Bathycoccus* genus was divided into two species, the polar and temperate *Bathycoccus prasinos* type B1 genome [13,22] and the tropical *Bathycoccus calidus* type B2 genome [21,23,24]. Thus the cosmopolitan nature of *Bathycoccus* from poles to equator is due to the combination of both B1 and B2 species.

In the bay of Banyuls, *Bathycoccus* and *Micromonas* bloom yearly from November to April, *B. prasinos* being one of the most abundant species [15,25]. The highly reproducible yearly occurrence of *B. prasinos* in the Banyuls bay during the last decade [15] raises the question of the persistence of a *B. prasinos* population adapted to the bay or of a variation of the population structure each year. In addition, since outside of the bloom period *Mamiellales* are virtually absent from the bay, is the *B. prasinos* bloom initiated by an uptake of resident “resting cells” in the sediment or by a fresh input carried by North western Mediterranean currents along the Gulf of Lion? At present no resting stages that can act as inoculum of subsequent blooms have been described for *B. prasinos*.

To assess the intraspecific diversity of *B. prasinos* worldwide and locally in the bay of Banyuls, we developed an efficient method to isolate strains together with whole genome sequencing by Oxford Nanopore Technology (ONT) in order to identify Structural Variations (SV) in *B. prasinos* genomes. Diversity markers designed from INDEL (insertion or deletion of bases in the genome of an organism) were used to genotype *B. prasinos* strains and populations from environmental samples.

## Methods

### Algal strains and culture conditions

World-wide *B. prasinos* strains were obtained from the Roscoff Culture Collection (RCC) centre (See Supplementary Table 1, Additional File 1). Strain RCC1105 from which the current *B. prasinos* reference genome originated [22] was lost and replaced by its clonal culture RCC4222. The strains were cultivated in 100 mL flasks in filtered artificial seawater (1×10^−3^ M TrisHCl pH 7.2, 24.55 g/L NaCl, 0.75 g/L KCl, 4.07 g/L MgCl2 6H2O, 1.47 g/L CaCl2 2H2O, 6.04 g/L MgSO4 7H2O, 0.21 g/L NaHCO3, 0.00138 g/L NaH2PO4, 0.075 g/L NaNO3, 1.28×10^−6^ g/L H2SeO3) supplemented with trace metals and vitamins B1 (300 nM) and B12 (1 nM). Cultures were maintained under constant gentle agitation in an orbital platform shaker (Heidoph shaker and mixer unima× 1010). Sunlight irradiation curves recreating realistic light regimes at a chosen latitude and period of the year were applied in temperature-controlled incubators (Panasonic MIR-154-PE).

### Cell isolation

Surface water was collected at 3 metre depth at SOLA buoy in Banyuls bay, North Western Mediterranean Sea, France (42°31′N, 03°11′E) approximately every week from December, 2018 to March, 2019 ; November, 2019 to March, 2020 and October, 2020 to April, 2021. Two ml aliquots were used to determine the quantity and size of phytoplankton by flow cytometry. For the 2018/2019 bloom, 50 ml were filtered through a 1.2-µm pore-size acrodisc (FP 30/1.2 CA-S cat N° 10462260 Whatman GE Healthcare Sciences) and used to inoculate 4 culture flasks with 10 ml of filtrate each. The sea water was supplemented by vitamins B1 and B12, NaH2PO4, NaNO3 and metal traces at the same final concentration as artificial sea water (ASW), antibiotics (Streptomycin sulfate and Penicillin at 50 μg/ml) were added to half of the cultures. The cultures were incubated under light and temperature conditions similar to those during the sampling date for 3-4 weeks. The presence of picophytoplankton was analysed by a BD Accuri C6 flow cytometer. Generally, superior results were obtained without antibiotics. Cultures containing at least 90% of picophytoplankton with only residual nanophytoplankton were used for plating on agarose (0.21%). Colonies appearing after 10 days were hand-picked and further cultured in 2 ml ASW in deepwell plates (Nunc, Perkin Elmer, Hessen, Germany) for 10 days. Cells were cryopreserved at this stage. (See Supplementary Figure 1, Additional File 2) Circa 500 clones were cryopreserved. At the same time, DNA extraction and PCR were performed in order to identify *B. prasinos* strains.

### DNA extraction, genome sequencing, assembly and PCR amplification

For PCR analysis, total DNA was extracted from 4 ml *B. prasinos* cell cultures according to the Plant DNAeasy Qiagen protocol. For whole genome sequencing by Oxford Nanopore Technology (ONT), DNA was extracted by a CTAB method from 100 ml culture principally based on Debladis et al. [26]. ONT libraries were prepared using the Rapid Barcoding Sequencing kit (SQK-RBK004) and deposited on R9.4 flow cells for sequencing run. For environmental samples, 5 litres of seawater at SOLA 3 metre depth were passed through 3 microns and 0.8 micron pore filters. DNA from cells collected on the 0.8 micron filters were extracted using the Plant DNAeasy Qiagen protocol with the addition of a proteinase K treatment in the AP1 buffer. PCR was performed using the Red Taq polymerase Master mix (VWR) with the required primers (See Supplementary Appendix 1, Additional File 3) and corresponding DNA. For sequencing, the PCR products were purified using the NucleoSpin Gel and PCR Clean-up kit (Macherey-Nagel reference 740609.50) and sent to GENEWIZ for Sanger sequencing.

Raw ONT Fast5 data were basecalled using Guppy 4.0.15 (https://nanoporetech.com) and the HAC model, and QC performed using NanoPlot 1.38.1 [27]. All reads with a QPHRED higher than 8 were retained and subjected to genome assembly using Flye 2.8 [28] under standard options for ONT data. Raw assemblies were then polished with 3 turns of Racon 1.4.3 with standard options [29] after mapping of raw reads on the previous round sequence using minimap2 2.24 (-ax map-ont mode) [30]. Final scaffolding was performed using Ragoo 1.1 [31] upon the original *B. prasinos* reference genome (GCA_002220235.1) [22]. Final QC of assemblies was performed using QUAST 5.0 [32].

### Variant calling and size control

Draft assemblies were aligned to *B. prasinos* reference genome using MUMmer 4.0.0 [33] nucmer (options: –maxmatch -c 500 -b 500 -l 100). Alignments were then filtered for identity (>90%) and length (>100bp) using delta-filter from MUMmer 4.0.0 (options: -m -i 90 -l 100). Variant calling on the resulting delta files was performed with SVanalyzer 0.36 (https://github.com/nhansen/SVanalyzer) SVrefine subcommand, and resulting VCFs were merged with bcftools merge 1.18 [34].

Size control on VCF was performed using a set of home-made Python scripts available at https://forge.ird.fr/diade/genomecodes under GPLv3 licence.

### Determination of Growth Rates

Cells isolated during December, 2018, January and February, 2019 in Banyuls bay were used in this experiment. For each culture condition the cell number was determined by flow cytometry daily, for 9 days. The growth rate was determined as Ln(N)/dT, where N is the cell concentration per ml and T the time (days). The determination of the maximal growth rate (μmax) and subsequent statistical analysis were conducted in accordance with the methodology described in Guyon et al. [35].

## Results

### INDELs diversity of selected *Bathycoccus prasinos* strains

With the aim of creating a genetic resource of natural variants for *B. prasinos*, we first undertook a search for genetic markers of diversity that could be used to differentiate isolates from natural communities (Fig. 1A). From the few strains available in the Roscoff culture collection, we examined the genetic diversity of *B. prasinos* and selected geographically dispersed strains, so that the markers could potentially be used locally as well as worldwide. Oxford Nanopore Technology (ONT) was used to sequence the genomes of 6 selected strains spread along a latitudinal gradient from the Baffin bay (67° N) to the Mediterranean Sea (40° N) (See Supplementary Table 1, Additional File 1). After individual *de novo* assembly, each genome was compared to the reference genome of the RCC1105 strain from the Banyuls bay [22] to identify 200 to 2,000 bp long INDELs. Selective amplification of these variable size INDELs by PCR generates different size DNA fragments that can be used to genotype new isolated strains as well as to track population diversity in environmental samples. Once sequenced, new genotypes further enrich the marker resource that could be used subsequently for genotyping *B. prasinos* isolates (Fig. 1A).

**Figure 1:**
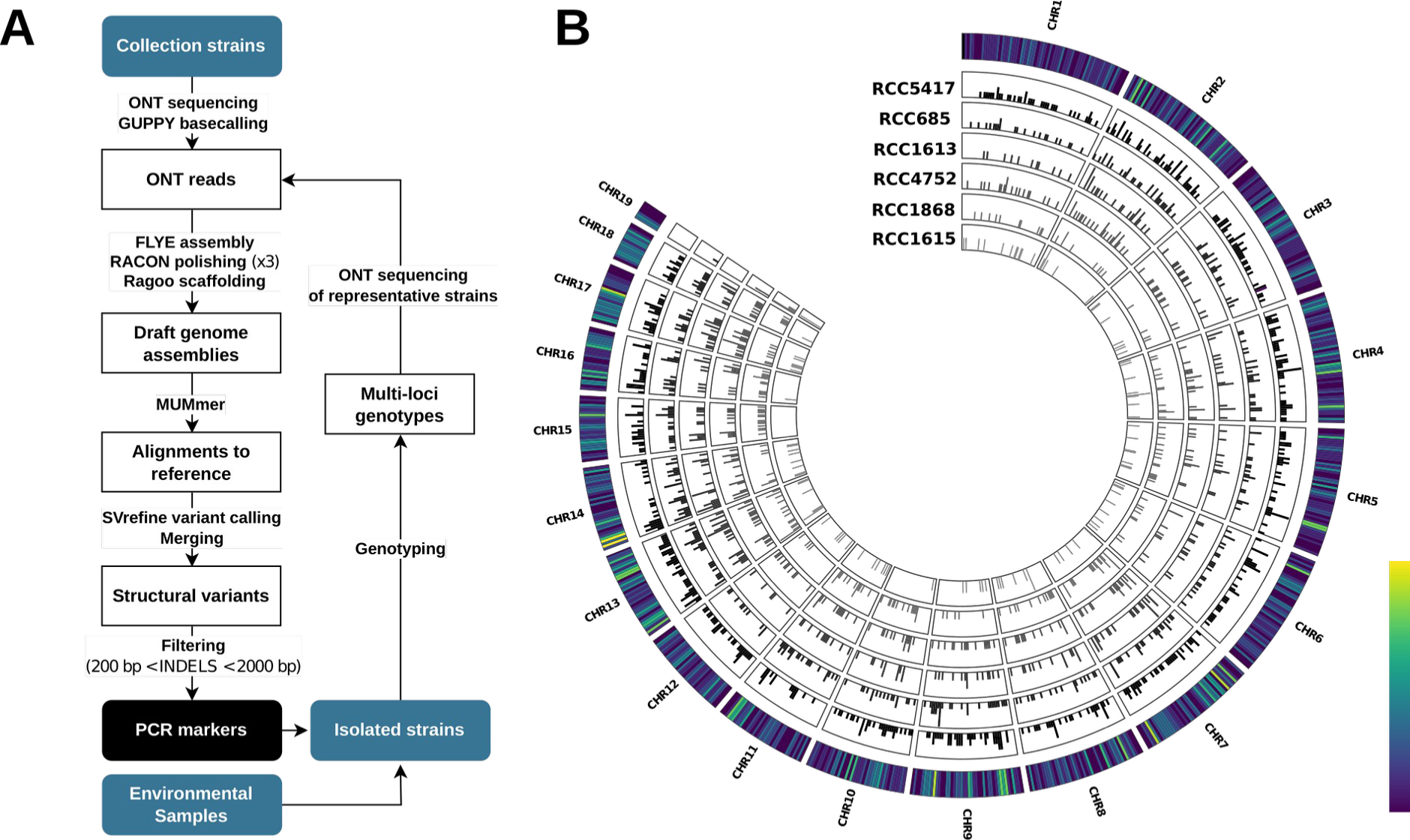
Structural variant identification in collection strains of B. prasinos. (A) Summary of the sequencing and analysis strategy for the identification of INDELs structural variations and the design of PCR diversity markers. (B) Distribution frequency of identified SV along a 20Kb sliding window on nuclear chromosomes. Each inner circle corresponds to the distribution of SV within a sequenced strain. The outer circle corresponds to the cumulative frequency of all sequenced strains.

A total of 1,346 unique SV were identified, the contribution of each strain being highly variable (Table 1). Strain RCC5417 from the Baffin bay was the most geographically distant from the reference strain, and also the most diverse with 536 SV of size > 200 bp identified. Conversely, RCC1615, with only a 4X sequencing depth, made the smallest contribution to SV identification up to 10,000 bp but it was the main contributor to larger SVs (>10,000 bp), with 73 SV detected. This may be due to the overall lower assembly contiguity and quality of this strain genome (Table 1; See Supplementary Table 1, Additional File 1). Despite these disparities, SV were evenly distributed along *B. prasinos* nuclear genome, with the exception of the outlier chromosome 19 [22], for which few variations were identified, most likely because of the hypervariable structure of this chromosome (Fig. 1B). In the case of the big outlier chromosome 14, no contrast in variant identification could be detected with other nuclear chromosomes.

**Table 1:**
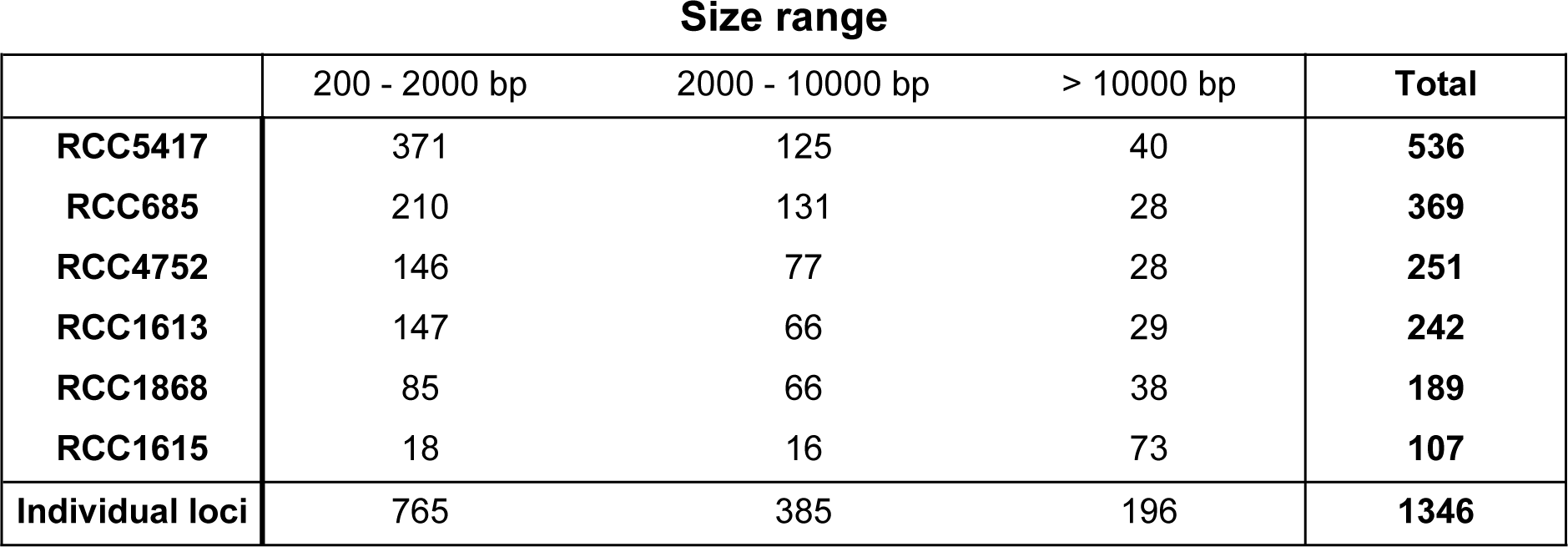
Number of structural variations identified in selected strains of *B. prasinos.* Individual loci corresponds to the cumulated number of variable genomic positions without duplicates.

|  | Size range |  |  | Total |
| --- | --- | --- | --- | --- |
|  | 200 - 2000 bp | 2000 - 10000 bp | > 10000 bp |  |
| **RCC5417** | 371 | 125 | 40 | **536** |
| **RCC685** | 210 | 131 | 28 | **369** |
| **RCC4752** | 146 | 77 | 28 | **251** |
| **RCC1613** | 147 | 66 | 29 | **242** |
| **RCC1868** | 85 | 66 | 38 | **189** |
| **RCC1615** | 18 | 16 | 73 | **107** |
| **Individual loci** | 765 | 385 | 196 | **1346** |

### Search for intraspecific diversity markers

To select putative PCR markers of *B. prasinos* diversity several criteria were considered. The size of the amplified fragment had to be between 0.2 and 2 Kb and sufficiently different among genomes to be unambiguously visualised on agarose gel after amplification with a single set of primers. In addition, selected INDEL SV had to be found in the genome of at least three strains. The aim of this selection was to identify the most divergent markers among the largest available genetic diversity of *B. prasinos*, with the expectation that some of these variations could potentially be found and tracked in local communities (Fig.1 A). In total, 765 unique SV with sizes ranging between 0.2 and 2 Kb were identified (Table 1). However, only 44 of them were found in at least three strains, thus meeting our criteria as putative markers (Fig. 2). Manual analysis of INDEL flanking sequences was performed to identify conserved sequences that could be used to design primers for PCR amplification of INDELS. Four markers were selected for experimental validation of INDELs by PCR (Fig. 3) (See Supplementary Appendix 2, Additional File 3).

**Figure 2:**
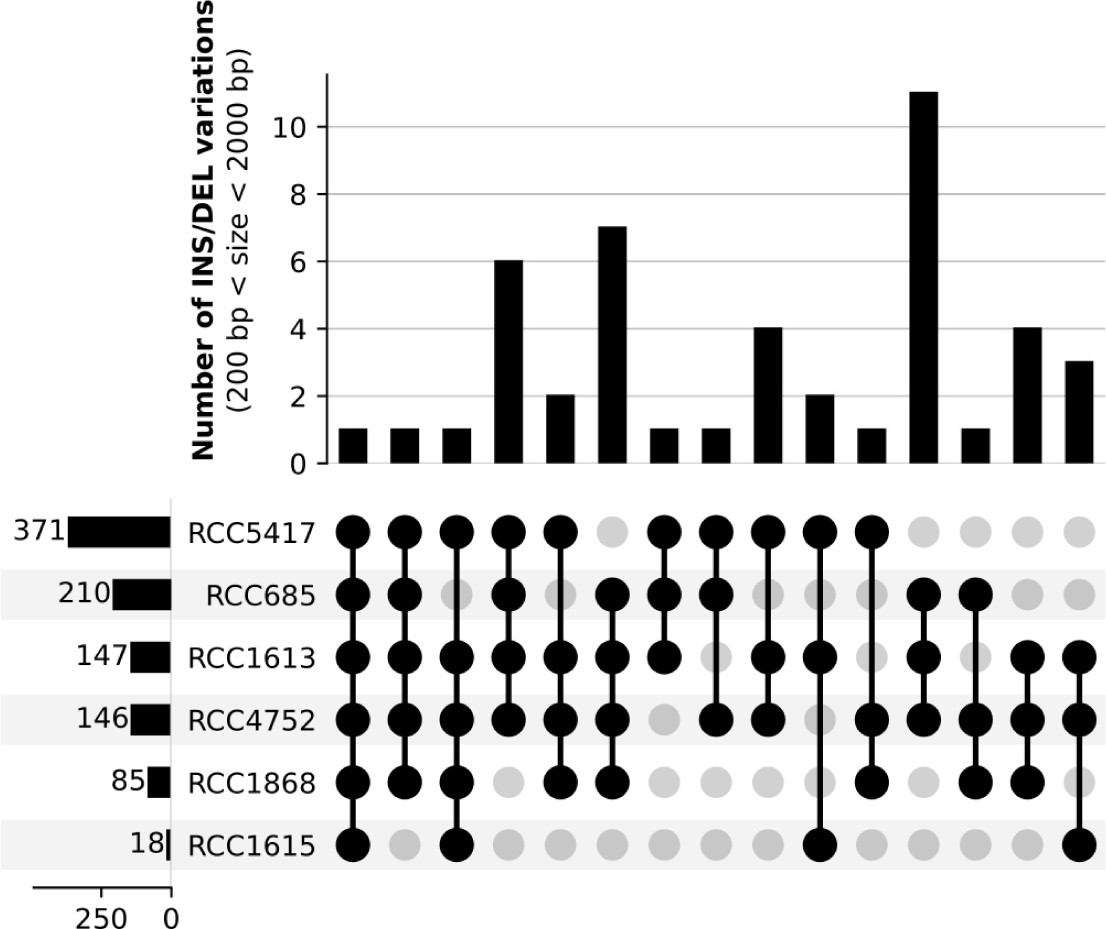
Distribution of structural variations shared by three or more strains. The vertical barplot represents the number of structural variations between 200 and 2000 bp detected only within the corresponding group of strains. The horizontal barplot represents the number of structural variations detected within the corresponding strain.

**Figure 3:**
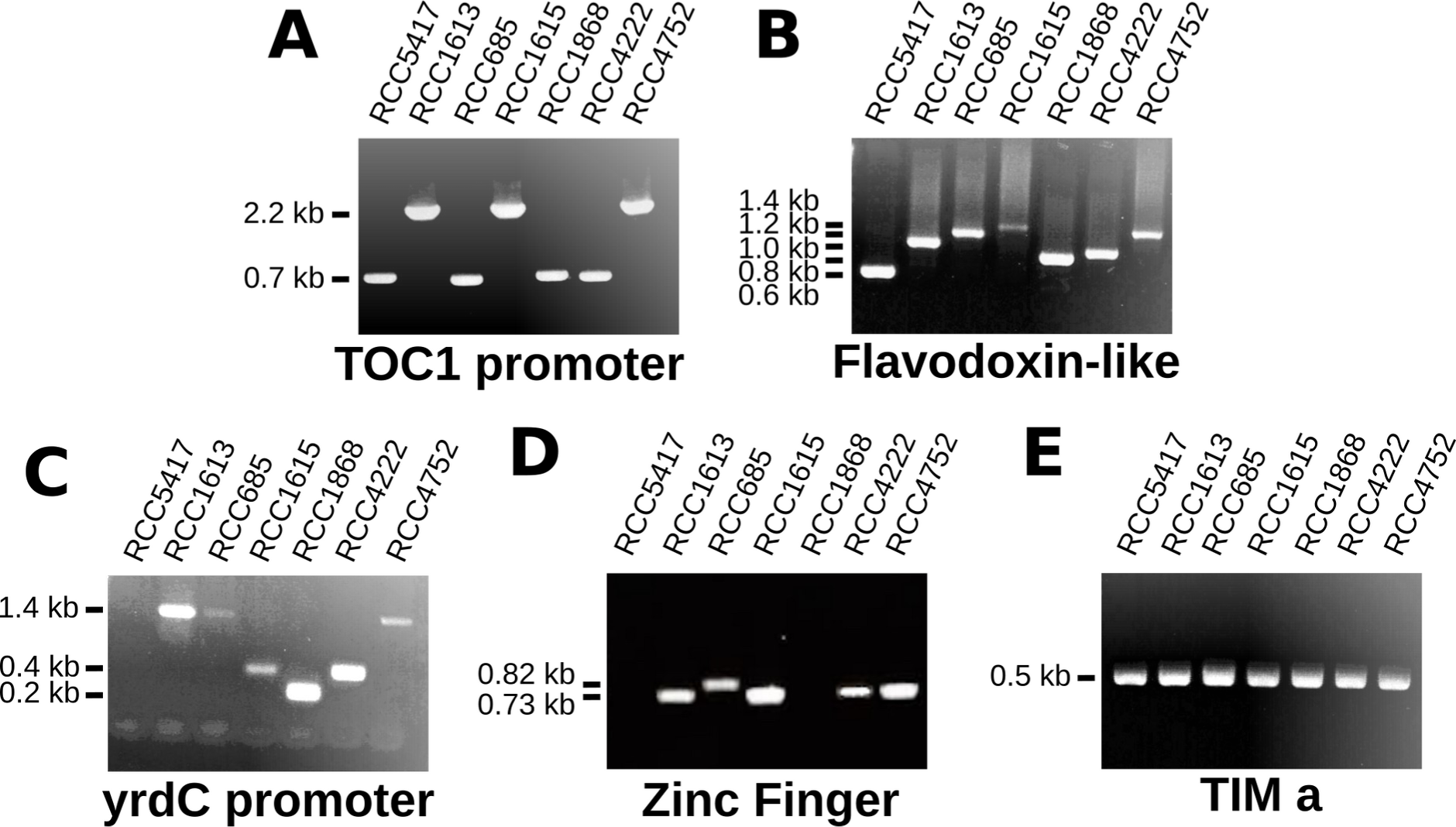
Diversity markers in collection strains. PCR amplification of diversity markers (A) TOC1 promoter, (B) Flavodoxin-like, (C) yrdC promoter, (D) Zinc Finger, (E) TIMa. Variations in amplification efficiency are due to mismatches between primer sets and genomes.

The “TOC1 promoter” marker, located on chromosome 17 (RCC1105: NC_023992.1:308225..308959) corresponds to an INDEL variation in the intergenic region upstream of gene Bathy17g01510, a homolog of circadian clock gene TOC1. This diversity marker is bi-allelic with amplified fragments of 700 or 2,200 bp in size (Fig. 3A).

The “Flavodoxin-like” marker, located on chromosome 3 (RCC1105: NC_024006.1:379834..380504) corresponds to an INDEL variation in the gene Bathy03g02080 which encodes a protein containing amino stretches repeats of various lengths. This marker shows 5 alleles producing amplified fragment lengths of 600, 800, 1000, 1,200 and 1,400 bp (Fig. 3B).

The “yrdC promoter” marker, located on chromosome 1 (RCC1105: NC_024008.1:800915..801320) corresponds to an INDEL variation in the promoter region of a yrdC domain-containing protein of unknown function (Bathy01g04300). In the tested strains, this diversity marker is tri-allelic and shows amplified fragments of 200, 400 or 1,400 bp.

Amplification efficiency was lower in strains RCC1615, RCC685 and RCC4752 and no amplification was obtained for strain RCC5417 (Fig. 3C).

Finally, the “Zinc finger” marker, located on chromosome 15 (RCC1105: NC_023994.1:446402..447132), corresponds to variation in the number of Zinc finger repeats encoded in the gene Bathy15g02320. This marker is biallelic and shows amplified fragments of 730 or 810 bp. No amplification was obtained for strain RCC5417 and RCC1868 (Fig.3 D).

The best sets of primers designed for markers “yrdC promoter” and “Zinc finger” show differences in amplification efficiencies between strains due to the lack of conservation in PCR primer sites in these strain genomes. However, the sequences of all amplified INDELs were verified by sequencing and correspond to the expected amplification.

A fifth additional marker, named “TIMa”, was designed to amplify a 530 bp fragment. This marker was selected because it is located in the BOC region of chromosome 14 at the border between a conserved region and an outlier region putatively involved in the definition of mating types in other *Bathycocacceae* (RCC1105: NC_023995.1:564926..565456) [19]. As expected this marker was amplified in the 6 genomes which would correspond to the same putative mating type (Fig 3E).

### Isolation and identification of *Bathycoccus prasinos* Multi Loci Genotypes in the Banyuls bay

Surface water was collected weekly at the SOLA buoy in Banyuls bay from December the 3^rd^, 2018 to March the 19^th^, 2019. During this period, the sea temperature ranged between 15.87°C in December and 10.68°C in February. We implemented a protocol to isolate *B. prasinos* cells based on filtration through 1.2 μm pore-size filters to remove larger cells of nanoplankton. This pore size allows the passage of *B. prasinos* cells (which size is estimated to be of 1.5 μm) and most importantly eliminates the potential larger predators. After a period of acclimation of two weeks in natural sea water supplemented with nitrate, phosphate and vitamins B1 and B12, cells were isolated by plating in low melting agarose supplemented sea water. Out of more than 400 colonies, only light green-yellow coloured colonies were picked and sub-cultured subsequently. In order to identify putative *B. prasinos* strains, amplification of a fragment of the LOV-HK gene (Bathy10g02360) was performed. These primers did not amplify LOV-HK sequences in other *Mamiellales* (*Ostreococcus* or *Micromonas*). The identity of these clones was further confirmed by ribotyping (amplification of a 2kb ribosomal DNA fragment followed by sequencing) (See Supplementary Appendix 1, Additional File 3). In total, 55 *B. prasinos* isolates were recovered for nine sampling dates (See Supplementary Table 2, Additional File 1).

The five markers, including the four diversity markers and the TIMa marker were used in combination to distinguish the different isolates of *B. prasinos* into Multi Loci Genotypes (MLG). Eight MLG were identified in Banyuls’ strains, MLG1 and MLG2 being dominant with 30 and 15 isolated strains respectively (Table 2). These two dominant MLG were distinguished only by the TIMa marker, which was amplified only in MLG1. For yrdC promoter marker, the 200 bp allele was dominant with an amplification in 54 isolates. The 1,400 bp allele, identified in worldwide strain, was not detected in the Banyuls bay strain. For the TOC1 promoter marker, the 1,200 bp allele was dominant with an amplification in 51 isolates. For the Flavodoxin-like marker, the 1,200 bp allele was dominant with an amplification in 49 isolates. From the 6 remaining isolates, 3 alleles could be identified, including a new 1,600 bp allele; however the 1,000 bp allele was not detected. Finally, for the Zinc finger marker, the dominant allele was the 730 bp one, amplified in 52 isolates.

**Table 2:**
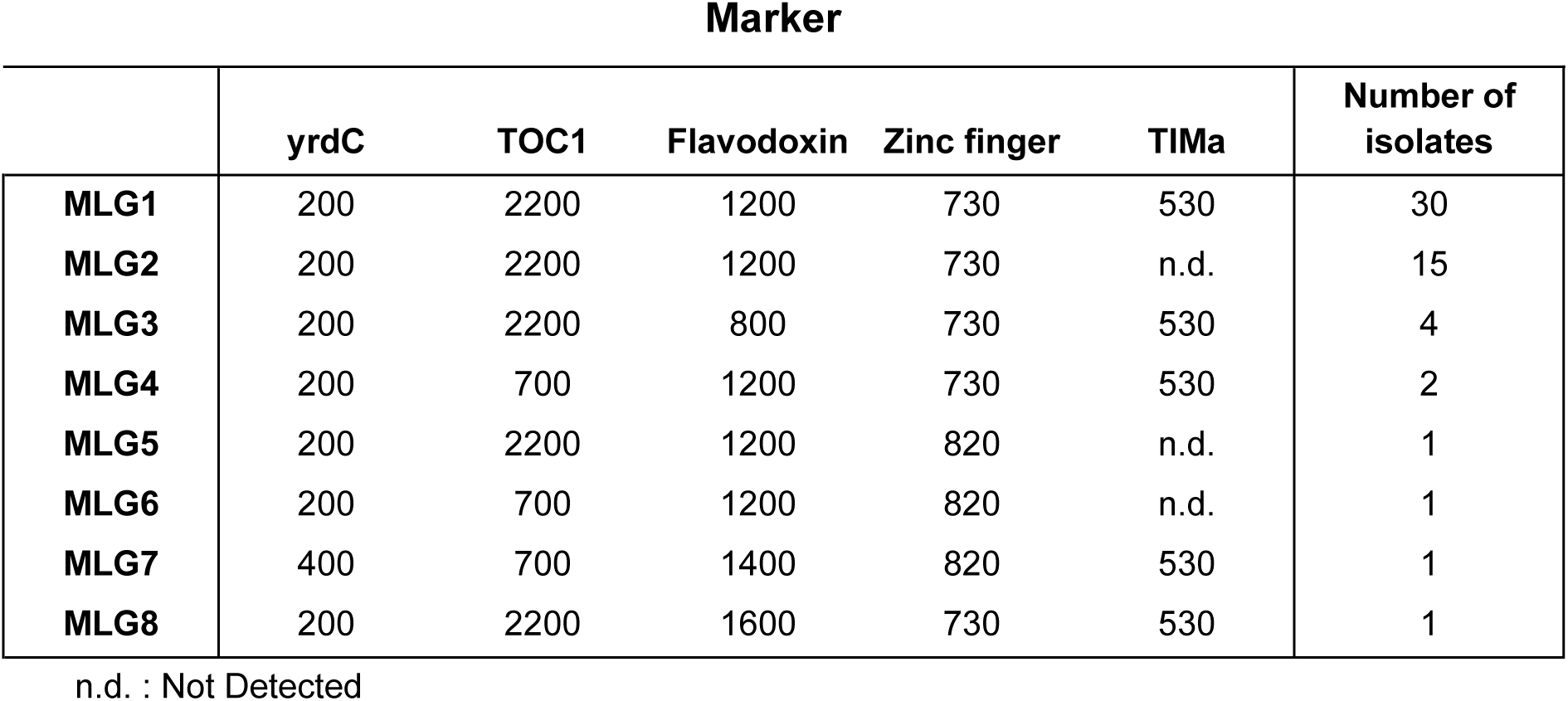
Marker characterisation of multi-loci genotypes in *B. prasinos* strains isolated from Banyuls bay. Marker values correspond to amplified sizes in base pairs.

The genotypic diversity observed by the MLG classification was also correlated with biological and physiological diversities of the strains. Indeed, by monitoring the growth response of representative strains under different temperature and light conditions corresponding to those observed at different parts of the bloom in the Banyuls bay, we showed clear differences in growth rate between MLG and conditions. Specifically, there were notable variations between strains from MLG1/MLG2 (isolated in December/January) and other tested MLG (isolated in February) (See Supplementary Figure 2, Additional File 2).

### Diversity of INDEL markers

Seven isolates corresponding to 6 MLG were selected for whole genome sequencing and assembly following the methodology described in Figure 1A. The genome assemblies revealed that the two MLG, which failed to amplify the TIMa marker, have a different haplotype of the chromosome 14 outlier region. A complementary marker called “TIMb’’ was designed from these genome assemblies, corresponding to a 1,300 bp amplified fragment. TIMb was detected in isolates for which TIMa was not amplified (See Supplementary Figure 3, Additional File 2).

Comparative analysis of the polymorphism in our markers between worldwide strains and newly isolated strains revealed that all five markers were at least biallelic in the Banyuls local population (Fig. 4A). Furthermore, new alleles were identified through the genotyping of Banyuls bay isolates for Flavodoxin-like and TIMa/b markers, thus extending the diversity observed of our markers. The pipeline of INDEL identification was applied to the seven strains sequenced from the Banyuls bay (Fig. 1A). This led to the detection of 211 unique INDELs of sizes ranging between 0.2 and 2 Kb (Fig. 4B). Within these loci, 79 corresponding to 37% of loci diversity in the Banyuls bay, were shared with worldwide strains (Fig. 4B). On average, each variant locus was found in 5.6 strains across the two populations (Fig. 4C).

**Figure 4:**
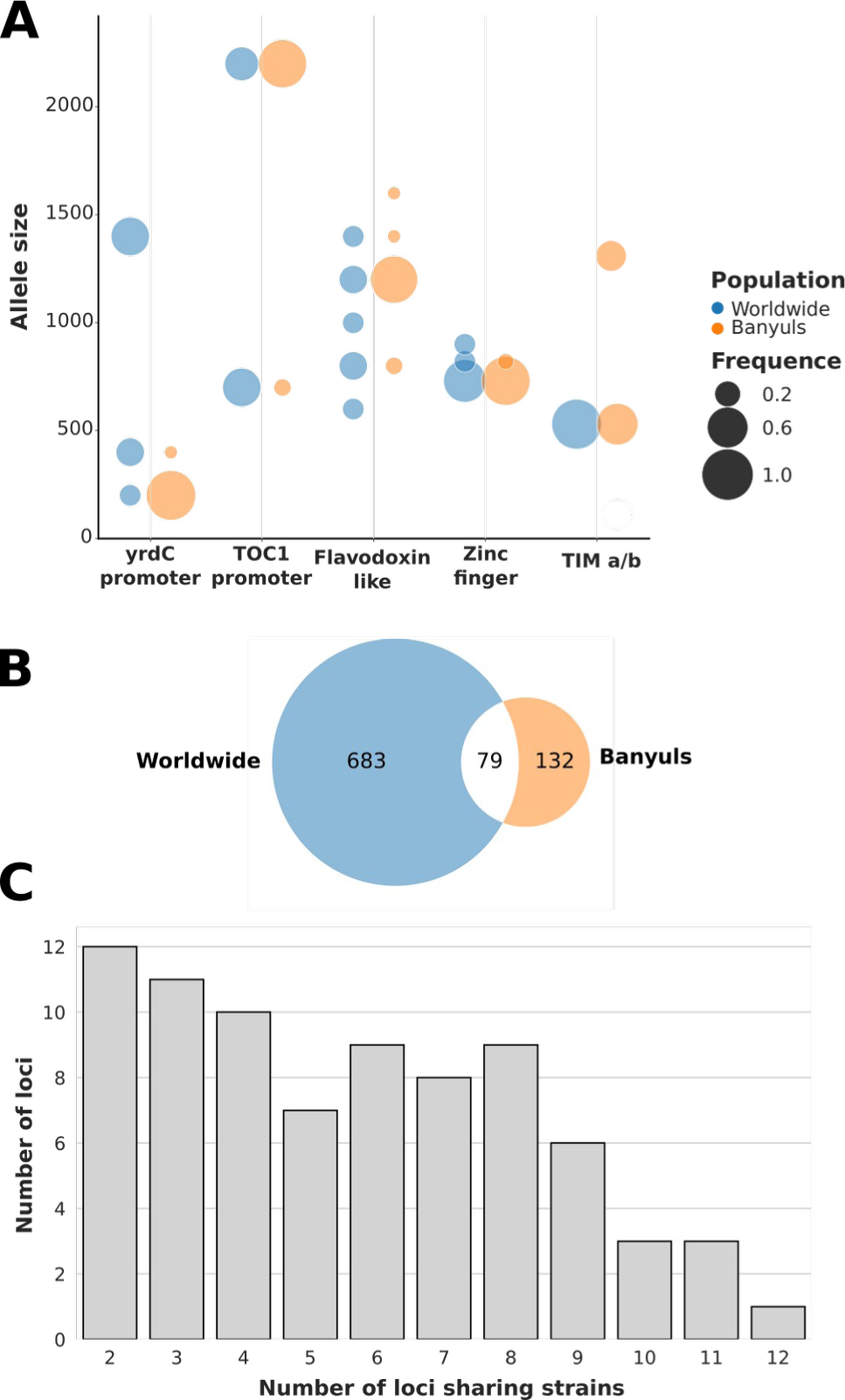
Worldwide and local diversity in marker alleles and variant loci. (A) Sizes and frequencies in marker alleles within worldwide strains (n=6) and local strains from the Banyuls Bay (n=55). (B) Specific and shared variant loci (200 < size < 2000 bp) between at least one worldwide sequenced strains (n=6) and one sequenced strains from the Banyuls Bay(n=7). (C) Distribution frequency of the 79 shared variant loci (200 < size < 2000 bp) in the 13 sequenced strains.

### Determination of allelic variants in environmental samples

The identification of dominant MLG during the 2018/2019 bloom raises the question of their yearly or occasional prevalence in the Banyuls bay. However, isolating strains is highly time consuming and not appropriate to follow population dynamics at higher frequency. To overcome this, our diversity markers were directly tracked on environmental DNA samples during 3 successive blooms from 2018 to 2021. Five litres of seawater were sampled once a week and filtered between 3 and 0.8 μm. DNA extracted from 0.8 μm filters was used for PCR analysis of presence/absence variations in alleles of yrdC promoter, TOC1 promoter and TIMa/TIMb markers (Table 3). Flavodoxin-like and Zinc finger markers could not be used on complex environmental DNA samples due to the presence of high background.

**Table 3:**
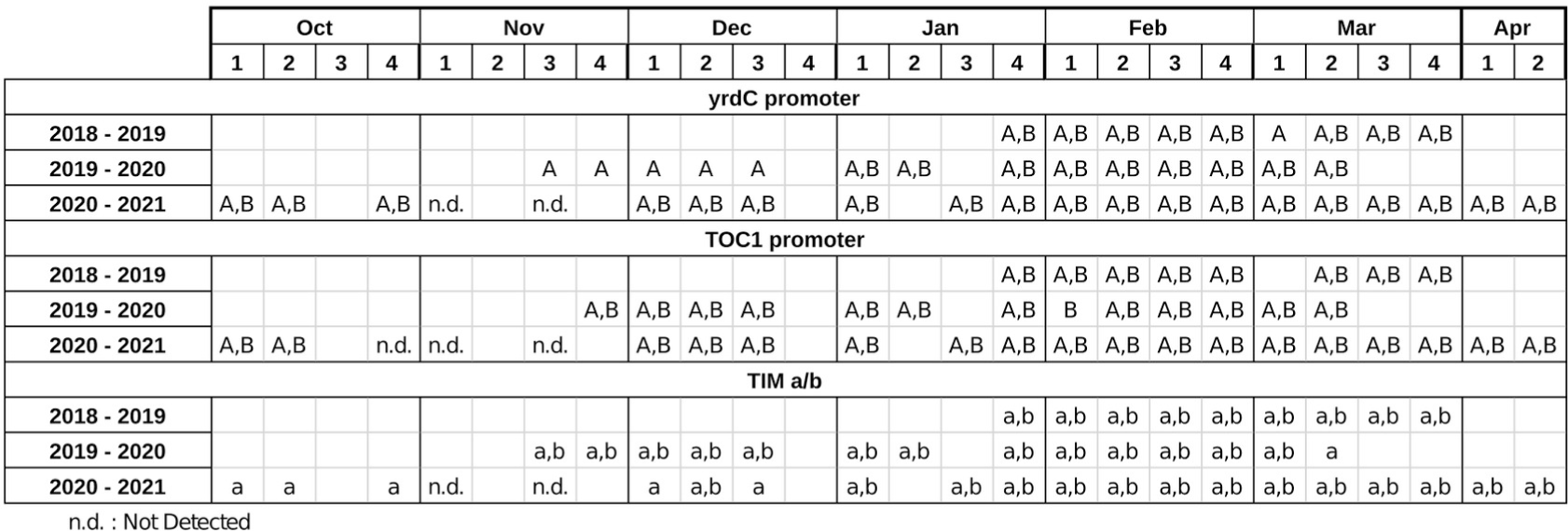
Marker alleles detection in environmental DNA samples from successives *B. prasinos* b looms in the Banyuls bay. YrdC promoter (A : 200bp, B : 400bp), TOC1 promoter (A : 2200bp, B: 700bp), TIM a/b (a : 530 bp, b : 1300bp). Missing datapoints are represented by empty cells.

The initiation of the 2019/2020 bloom was defined by the exclusive presence of the 200 bp allele of yrdC promoter marker from the third week of November to the third week of December. This was followed from January to March by the unambiguous detection of both the 200 bp and the 400 bp alleles. During this bloom, TIMa and TIMb markers were both amplified in all samples, with the exception of the second week of March, during which only the TIMa marker was detected.

Several distinctions can be done with the 2020/2021 bloom, the first one being that no amplification was detected in November samples, an observation consistent with the overall decrease in picophytoplankton abundance (See Supplementary Figure 4, Additional File 2). For yrdC promoter marker, the 400 bp allele was observed in October and December, hence between one and three months earlier than in the 2019/2020 bloom. In addition, only TIMa was amplified in October and December 2020. Both alleles of the TOC1 promoter marker were amplified in nearly all samples where *B. prasinos* could be detected and were therefore not informative in this context (Table 3). From the study of these successive blooms, we were able to conclude that their onset differed in both chronology and population diversity. Furthermore, we demonstrated that population diversity changes both within and between blooms at a given site and that diversity markers, designed from the worldwide diversity, could be used to successfully follow these local population dynamics.

## Discussion

We aimed to develop a time and cost effective method to produce a biological resource and characterise the intraspecific diversity of the marine microalga *B. prasinos*. In recent years, the cost of high-throughput sequencing has fallen sharply, favouring blind sequencing approaches without prior genotyping [19]. However, the isolation of new strains from complex microbial communities is time consuming, requiring multiple steps from sample isolation in remote areas of the ocean, to incubations, shipment and isolation by subculturing on agarose plates. Moreover, the incubation step can lead to the amplification of strains that, without genotyping, could be sequenced multiple times, while missing rare genotypes that contribute to the fitness of a species under changing environmental conditions. Given the bottleneck steps of strain isolation and whole genome sequencing in complex microbial communities, we turned to PCR amplification as a potentially effective approach for (i) preliminary screening of environmental samples to retain and incubate only those containing *B. prasinos* cells, (ii) identifying incubates containing *B. prasinos* for subsequent isolation of strains, (iii) genotyping and identifying the main MLGs prior to sequencing. These first steps of screening by PCR could be potentially performed during oceanographic cruise to guide and optimise sample selection and culturing on boats. Remarkably, our protocol proved efficient and selective for *B. prasinos* isolation since *Ostreococcus* and *Micromonas* strains were not obtained even though *Micromonas* is as abundant as *B. prasinos* in the Banyuls bay in winter [15]. This is all the more important as flow cytometry cannot distinguish the *Bathycoccus* genus from other *Mamiellales*.

### Design of diversity markers from INDELs structural variations

While PCR, combined with DNA fragment size estimation by gel electrophoresis, has already been used to follow intraspecific diversity [12], this approach requires the specific amplification of size-varying loci. This leads to challenges such as identifying putative marker loci that vary in size, are sufficiently conserved to be representative of subpopulations and can be found in conserved regions for which PCR primers can be designed. To address this, we explored the utility of INDEL variations within coding sequences, which offer higher stability in clonal populations compared to SNP and microsatellite markers, and are therefore good candidates for the design of size markers to track intraspecific diversity [36,37]. Furthermore, information on such INDELs variation and on repeated/low complexity sequences are now becoming easily available through the leveraging of the recent ONT and PacBio long read sequencing technologies, providing access to structural variations and marker designs previously overlooked [31,38,39].

By long read sequencing a selection of only six geographically distinct strains available in culture collection, we were able to obtain a first picture of the genetic diversity of *B. prasinos*. T he implemented genome assembly pipeline provided the draft genomes required for comparison and identification of structural variations. This resulted in the initial identification of 683 INDEL variation candidates, including 44 meeting our criteria for the design of diversity markers. While each candidate marker must be validated experimentally, this demonstrates the effectiveness of long reads whole genome sequencing for the identification of diversity marker loci using a limited biological resource. Four loci were selected for PCR validation, resulting in the diversity markers Flavodoxin-like, TOC1 promoter, yrdC promoter and Zinc finger. All markers were successfully amplified in a majority of strains even though a lack of conservation in PCR primer sites resulted in uneven amplification signals for yrdC promoter and Zinc finger markers. As such, nucleotide diversity remains a limiting factor for the design of universal markers when considering phylogenetically distant strains. Nonetheless, the pipeline developed in this study, by systematically mapping all INDEL and iteratively integrating them in the database, provides an extensive resource to identify novel markers (Fig. 1a; Fig. 4).

As expected, fewer alleles were detected in the local Mediterranean population for the designed markers, but genetic diversity was still observed for all of them. Since all markers correspond to structural variations either within coding sequences or regulatory sequences, this raises the question of whether they play a role in local adaptation. For instance, in the case of the central circadian clock TOC1, the variation was found in the promoter which contains a key *cis*-regulatory Evening element that is essential for the circadian regulation of TOC1 [40]. However, no clear pattern of allele distribution was identified neither in the worldwide strains nor in the sampled Banyuls population. The potential role of these INDELs in adaptation would therefore require additional experimental investigation.

### Genotyping and phenotyping in local populations of *B. prasinos*

The five diversity markers developed in this study were tested in combination on 55 freshly isolated Mediterranean strains of *B. prasinos*. This resulted in their classification into eight multi-loci genotypes. This highlights, firstly, the efficiency of our protocol for isolating *B. prasinos* strains from complex seawater samples and its effectiveness in preserving genotypic diversity. This also includes the TIMa/b markers which were manually designed from assembled genomes to detect different haplotypes of the big outlier chromosome 14 (BOC) as described by Blanc-Mathieu et al. [19] in *Ostreococcus tauri*. This marker combination allowed us to confirm the existence of at least two distinct haplotypes of the BOC in the *B. prasinos* subpopulation from the Banyuls bay, and thus can be used to distinguish these putative mating types in newly isolated strains and environmental samples.

A high level of intraspecific diversity has been reported in the few studies describing the genetic diversity of blooming phytoplankton populations [41–44]. For example, in diatoms (although not directly comparable as microsatellite markers have a higher mutation rate than other regions of the genome), more than 600 individuals were genotyped using microsatellite markers, and it was estimated that the blooming population was comprised of at least 2400 different genotypes [41]. Therefore, the observation of eight MLG in *B. prasinos* mediterranean population is most likely an underestimation due to the low number of markers used and their limited polymorphism. However, further improvement in the accuracy of our marker set can be achieved in an iterative way: through the identification of new size alleles (as exemplified by the 1,600 bp allele of the flavodoxin-like marker identified in the Banyuls population), or by the sequencing and integration of newly isolated strains distinguished by our original set of primer, further increasing the number of candidate loci. In this study, the integration of 7 newly sequenced genomes from the Banyuls bay nearly doubled the number of putative universal markers and provided population specific variants that could be used in local subpopulations tracking.

The first physiological characterization of selected strains revealed distinct growth responses, the strains isolated in December and January showing reduced growth under lower temperature (MLG1 and MLG2) and short photoperiod (MLG1) which suggest that these strains may correspond to seasonal ecotypes. Before concluding whether the different MLG classes correspond to different ecotypes, it will be necessary to characterise the physiological response of more strains, in particular those belonging to each of the dominant MLG1 and MLG2 classes.

### Tracking intraspecific diversity directly in environmental samples

Isolation of *B. prasinos* strains is too laborious and costly to be performed systematically on a large scale. For this reason, we attempted to assess intraspecific diversity in environmental samples during three consecutives annual blooms of *B. prasinos* in the Banyuls bay. Although flavodoxin-like was the most resolutive marker in isolated strains, PCR on environmental DNA yielded a smear on electrophoresis gel, possibly due to high polymorphism. Three diversity markers, TIMa/b, yrdC and TOC1 promoter, were suitable for amplifying environmental DNA, highlighting local population dynamics within and between bloom events. While this approach based on PCR amplification provides only qualitative data at this stage, approaches based on quantitative PCR could be developed to track the dynamics of genetic diversity in *B. prasinos* blooms directly from environmental samples [45]. As such, by accessing a level of structural diversity hardly obtainable through metagenomic short read mapping, it is complementary with approaches of metagenomic characterisation through SNP, as proposed by Da silva et al. [46].

Variations in interannual presence/absence of alleles raise the question of the nature of the highly reproducible yearly occurrence of *B. prasinos* in the Banyuls bay. Seasonal blooms may result either from re-activation of “dormant/survivor” cells from the water column (whose genetic fingerprint will determine the genetic profile of the next bloom) or by yearly *de novo* seeding by cells carried by the north Mediterranean current along the Gulf of Lion. At first glance, our preliminary qualitative results are in favour of the introduction of a new population rather than “resurrection” of cells from the previous bloom, since the allelic composition is distinct between the end of a bloom and the onset of the next (Table 3). By monitoring the temporal population structures of the dinoflagellate *Alexandrium minutum* in two estuaries in France, Dia et al. [44] showed that interannual genetic differentiation was greater than intra-bloom differentiation. Alternation of genotypes/populations has also been observed with diatoms in the dominance of one of the two sympatric populations of *Pseudonitzschia multistriata*, which could be due either to environmental factors favouring one population over the other or intrinsic factors coupled to the obligate sexual life cycle of *P. multistriata* [47]. Thus the observed fluctuations in allele frequencies could equally be the result of new inoculum from currents or sexual reproduction. Even though sexual reproduction has not been demonstrated in *B. prasinos*, there is genomic evidence that it may occur [48], and potential mating types were identified in our study (TIMa/TIMb markers). Sexual recombination generates new combinations of alleles, whereas clonality favours the spread of the fittest genotype through the entire population [44]. Erdner et al. [49] propose for *A. fundyense* that mitosis is the primary mode of multiplication during blooms, whereas mating is triggered presumably in response to unfavourable conditions at the end of blooms, with vegetative cells not overwintering in the water column. Knowing that (i) *B. prasinos* blooms are followed by severe bottlenecks between one bloom and the next [15], (ii) allele occurrences were different between the end of one bloom and the onset of the next, and (iii) structural markers are very stable in mitotic dividing cells, the hypothesis of rare vegetative cells remaining in the water column between the blooms is unlikely, except if those remaining cells were produced by sexual reproduction. The alternative hypothesis of new strains brought by current is however still equally probable. To test these hypotheses, more quantitative approaches based on real time quantitative PCR, or direct mapping of INDEL markers on the 7 year metagenomic time series in the Banyuls bay could potentially be used [50].

## Conclusions

In this paper, we describe the development of a new type of diversity markers based on INDEL structural variants obtained by long read sequencing. After validation on freshly isolated individuals, markers were used *in situ* on environmental samples. Whole genome sequencing of the new MLG refined a marker resource that could be further used to guide strain isolation and genotype new strains from the world ocean. In addition, the assessment of the physiological performances of genotyped strains suggest the presence of seasonal ecotypes in the Banyuls bay. Finally, PCR on environmental samples provided insight into the genetic diversity of *B. prasinos* seasonal blooms.

This INDEL marker resource represents a new tool for grasping the maximum diversity of newly isolated *B. prasinos* with the long term goal of identifying putative molecular mechanisms involved in adaptation to environmental niches in this cosmopolitan genus. The marker resource could also potentially be used to check culture purity or to follow specific genotypes in competition experiments in culture or in microcosms. The developed pipeline aimed at optimising sequencing efforts and building a collection of sequenced genomes representative of *B. prasinos* intraspecific diversity. This pipeline could easily be applied to other cosmopolitan and cultivable planktonic species that are available in collection, whether they are related to *B. prasinos* (e.g. *Ostreococcus* RCC809, *Micromonas commoda*, *Micromonas pusilla*) or share similar genome features.

## List of abbreviations

ASW: Artificial Sea Water
BOC: Big Outlier Chromosome
INDEL: Insertion/Deletion
ONT: Oxford Nanopore Tech
PCR: Polymerase Chain Reaction
RCC: Roscoff Culture Collection
SV: Structural Variation

## Declarations

### Availability of data and materials

Whole genome sequences and basecalled reads of the strains sequenced in this study as well as variant calling outputs are available at Zenodo https://doi.org/10.5281/zenodo.11395682. Strains isolated in Banyuls are available on request. All codes are available at https://forge.ird.fr/diade/genomecodes under the GPLv3 licence.

## Funding

The work was financed by an internal LOMIC Microproject to MD and the ANR Clima-Clock, ANR-20-CE20-0024 to FYB.

## Author contribution

MD, FYB and FS conceived the work and acquired funding. MD extracted high molecular weight genomic DNA. CM and MD performed the ONT sequencing. LD and FS performed bioinformatic analysis. MD designed and validated the diversity markers. PS analysed the seawater samples by flow cytometry. MD and VV isolated Banyuls strains during the winter bloom, genotyped by MD and JCL. MD and JCL determined the diversity of seawater samples. MD, LD and FYB wrote the article. JCL and FS took part in the critical reviews. All authors approved the final version for submission.

## Acknowledgments

We are grateful to the captain and the crew of the RV ‘Nereis II’ for their help in acquiring the samples. We thank the Roscoff Culture Collection for providing access to collected strains. Additional ONT were performed with the help of Christel Llauro and Marie Mirouze LGDP. The authors acknowledge the ISO 9001 certified IRD itrop HPC (member of the South Green Platform) at IRD Montpellier for providing HPC resources that have contributed to the research results reported within this paper (URL: https://bioinfo.ird.fr/ - http://www.southgreen.fr). We thank Thomas Roscoe for critically reading the manuscript.

## Notes

### Competing Interest Statement

The authors have declared no competing interest.

### Summary of Updates

This version has been revised in order to provide more technical/methodological aspects to the paper, and to strengthen its message. The original goal, ie having a robust method for phytoplancton genotyping, has been assumed all along the manuscript

