## Additional File 2 for "An INDEL genomic approach to explore population diversity of phytoplankton"

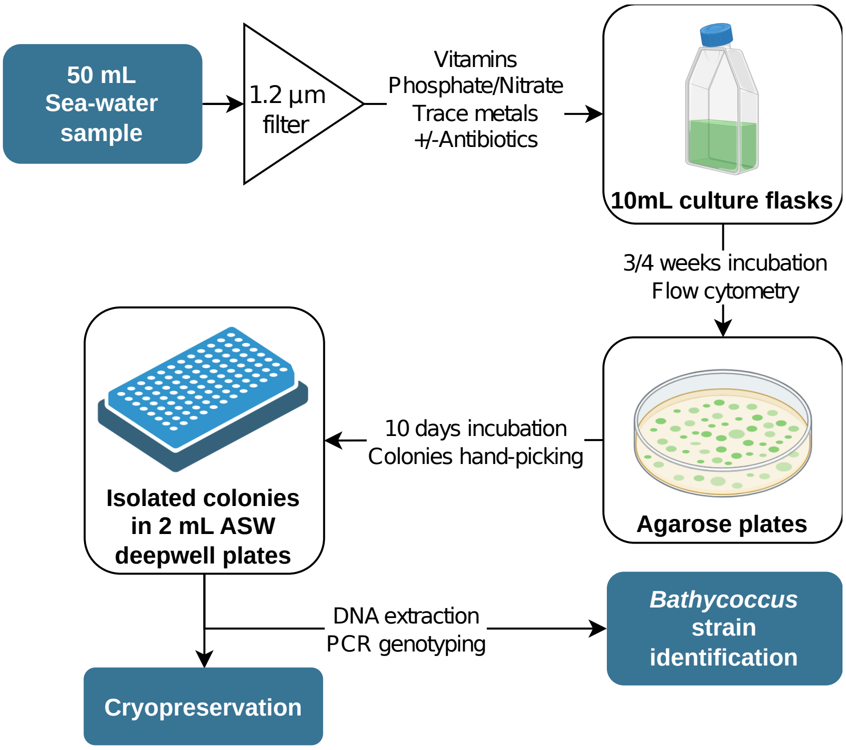


**Supplementary figure 1 : *B. prasinos* cells isolation protocol**
Graphical summary of the method for isolating *B. prasinos* cells from sea-water samples.

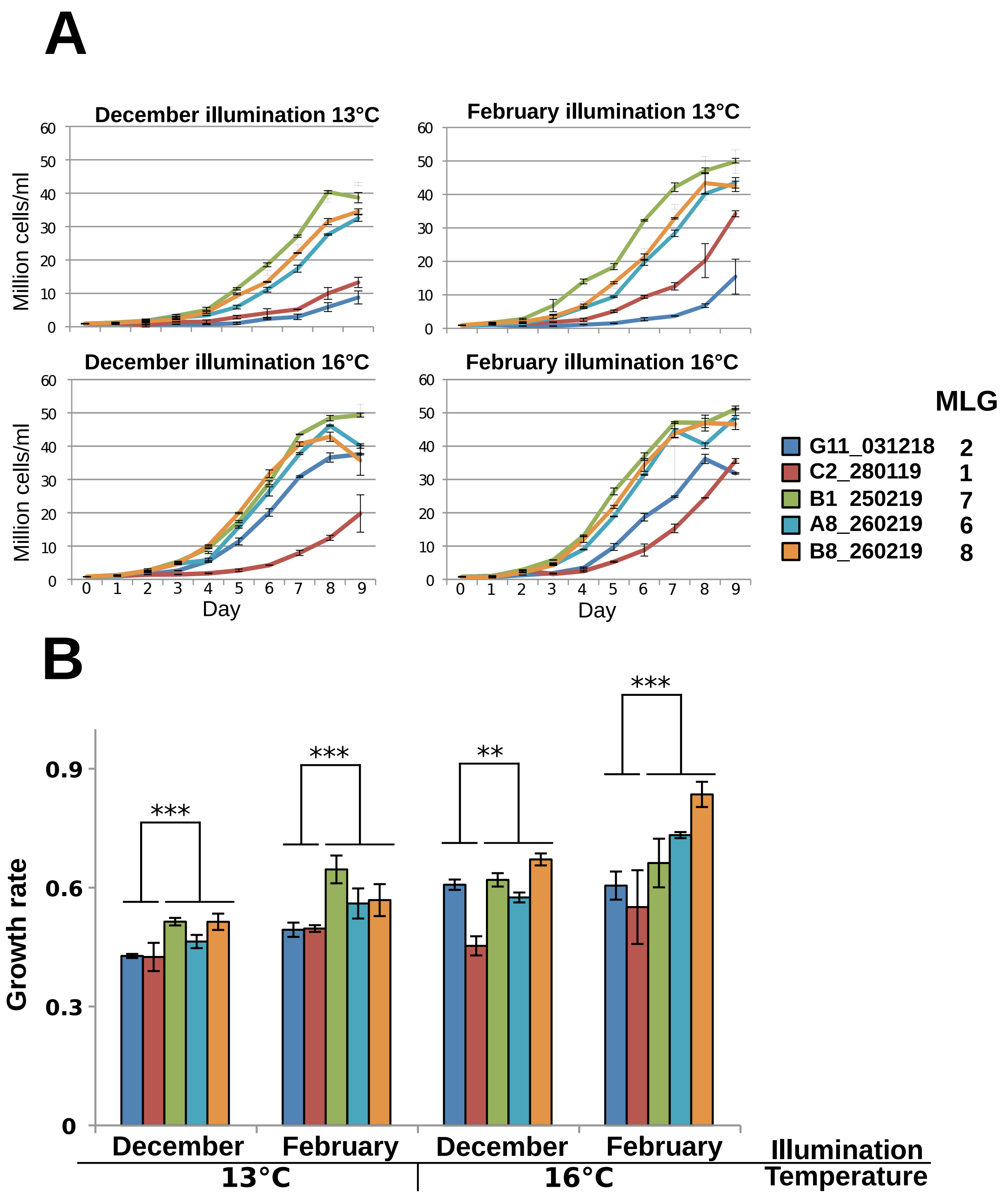


**Supplementary figure 2: Growth rate and curves of *B. prasinos* Banyuls isolates**

(A) The growth curves and (B) growth rates of five *B. prasinos* strains isolated from the Banyuls bay were determined under four different conditions of illumination and temperature by daily sampling for 9 days. Cell concentration was determined by flow cytometry and is expressed as $10^6$ cells/ml. Error bars correspond to standard deviation. One way analysis of t-test variance was used to compare strains growth rate in each condition.

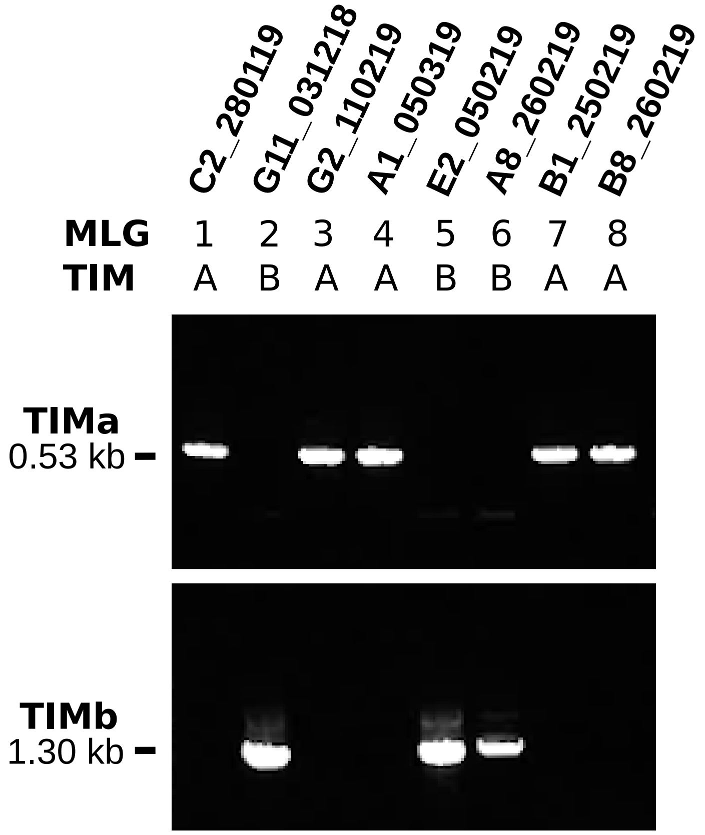


**Supplementary figure 3 : TIM a/b marker diversity in isolated strains**
PCR amplification using TIM a primers (top) and TIM b primer (bottom) in isolated strains from the Banyuls Bay representative of the 8 MLG.

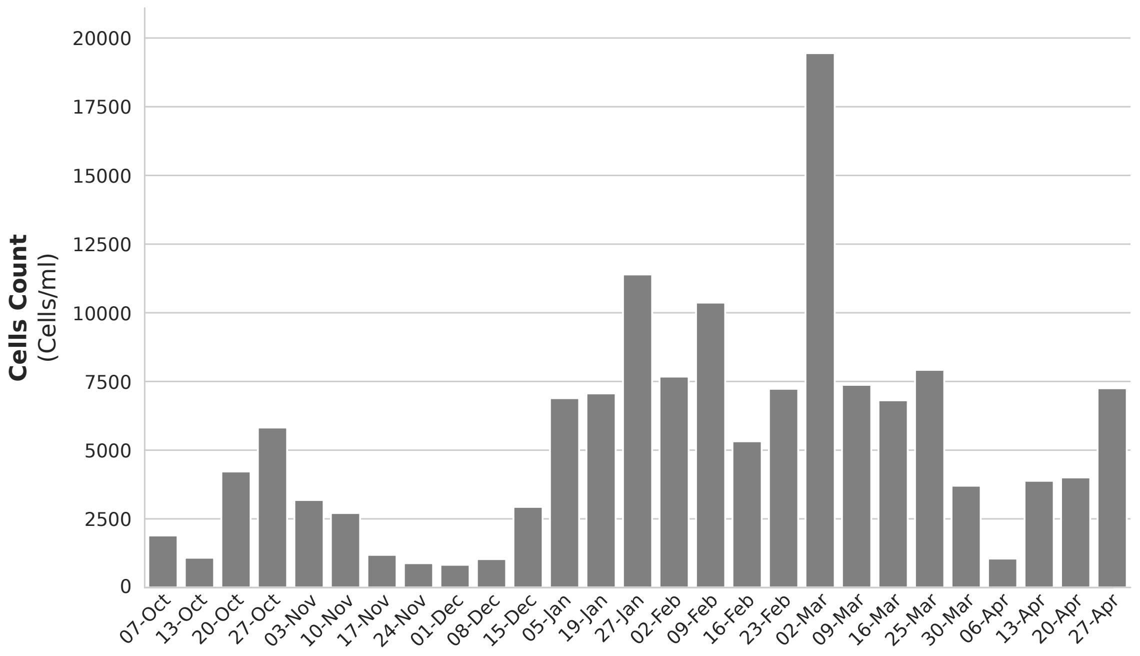


**Supplementary figure 4: Eukaryotic picophytoplankton abundance during the 2020/2021 winter bloom.** Flow cytometer counts of picophytoplankton eukaryotic cells in seawater sampled weekly between October 2020 and April 2021 at the SOLA buoy in the Banyuls Bay.
