## Additional File 3 for "An INDEL genomic approach to explore population diversity of phytoplankton"

### Appendix 1 : Primers used in this study

#### 1. Identification of Bathycoccus cells

a) Differences in the sequence of the LOV-histidine kinase (Bathy10g02360) gene have been used to identify *B. prasinos* cells.

BATHYK2F : GCCGTGATATCGTATCAAGTGTCC
BATHYK2R : CGAGAGCATTCAGCTTTTCCTCGA

b) Two kb of 18S ribosomal RNA was amplified by universal primers Euk 328f and Euk 329r and sequenced with Euk 528f

Euk 328f : ACCTGGTTGATCCTGCCAG
Euk 329r : TGATCCTTCYGCAGGTTCAC
Euk 528f : GCGGTAATTCCAGCTCCAA

#### 2. Diversity marker yrdC promoter

Variation in the promoter region of Bathy01g04300, a yrdC domain-containing protein.

MDB33 and MDB34 amplify the promoter of yrdC in all strains. MDB33 and MDB35 amplify a fragment only in the strain with an additional ANK repeat coding gene.

MDB33 : TACCGCAGCACGACTGCGGT
MDB34 : TTAATAAAGTGGTGGATTAACCTC
MDB35 : AGTTCCTCGCACTTCCTCTTCA

#### 3. Diversity marker TOC1 promoter

Primers MDB7 and MDB9 are used to amplify part of the TOC1 (Bathy17g01510) promoter region. MDB11 and MDB12 in strains containing an insertion into TOC1 promoter

MDB7 : ACTTGCTCGTTTCGAAAGTCGAA
MDB9 : TTCGAGAACGACGACGATCTCT
MDB11 : AGAGTTTGGAGGAGCCTCTGAA
MDB12 : ATACTCTTGGACCGTTTCCTCT

#### 4. Diversity marker Zinc finger

Variation in the number of repeats in the zinc finger protein (C2H2 Family) encoded by Bathy15g02300. MDB68 and MDB69 amplify a 730 bp DNA fragment from the reference genome and 820 bp in variant genomes.

MDB68 : CTAACGATGAGTACGCATGTGG
MDB69 : TGAGCAGCCGGCGGAAGAATC

#### 5. Diversity marker Flavodoxin-like

Variation in the number of repeats in the flavodoxin-like protein encoded by gene Bathy03g02080.

MDB40 : GAGAGAAGATTGAGGCGGAACA
MDB41 : TTCGGCAGCTGCTTTCGCTTCA

#### 6. Diversity marker TIM a/b

Variation of the C ter half of the protein encoded by Bathy14g03100.
With MDB57 and MDB58 amplification of a 530 bp band in reference strain.
With MDB57 and MDB59, a 1.3 Kb band in variant isolates.

MDB57 : GTGCAACTAATGTGCAGGAGCA
MDB58 : TTTCGACCACACGACGCACACT
MDB59 : TCTACGTGACCTGCATTGACTA

### Appendix 2 : Detailed description of diversity markers

#### A. Marker yrdC promoter

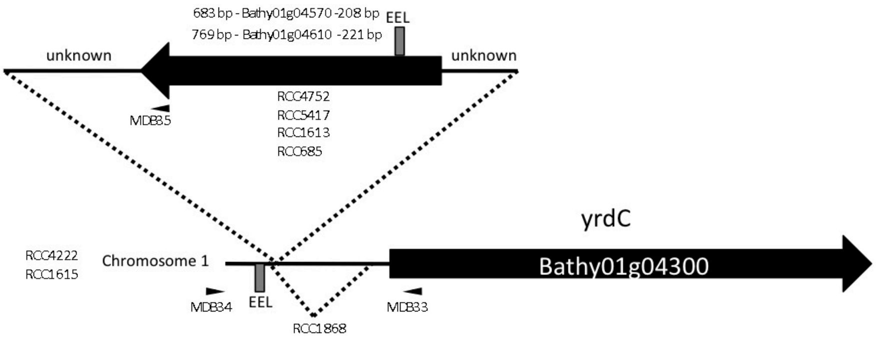


**Organisation of the *yrdC* promoter region on chromosome 1**
Schematic representation of the upstream region of gene Bathy01g04300 encoding an homolog the bacterial *yrdC* protein. Most strains possess a large insertion encoding a protein with Ankyrin (ANK) repeats and an additional Evening Element like (EEL) *cis*-element (grey box). In RCC1868, a deletion of 200 bp positions the EEL box closer to the putative start of transcription of Bathy01g04300. The position of the primers used for amplification and sequencing is indicated by arrowheads. Dotted lines indicate insertion or deletion.

Bathy01g04300 encodes a yrdC domain-containing protein of unknown function. In *Escherichia coli*, yrdC binds preferentially to double-stranded RNA, consistent with a role of the protein in translation (Teplova et al. 2000). A variable organisation was identified in the promoter region of Bathy01g04300 in comparison to the reference genome RCC1105/RCC4222. Only RCC1615 shares an organisation similar to that of RCC4222. In RCC1868, no insertion but a deletion of 212 bp was found. In other strains, a gene encoding a protein of unknown function possessing ANK repeats similar to the gene products of Bathy01g04610 (68% amino acid identity), Bathy01g04570 (67%) and Bathy11g02720 (57%) was inserted between an Evening Element-like (EEL) *cis* element and the start of Bathy01g04300. EEL boxes are recognised by Myb transcription factors CCA1 (Bathy05g02420) (http://plantregmap.cbi.pku.edu.cn/), regulators of the circadian rhythm. This arrangement was found in RCC5417, RCC1613, RC685 and RCC4752. The inserted gene should be considered as a putative ANK repeat additional gene, a family containing 186 genes in RCC1105 (Moreau et al. 2012).

##### Multiple nucleotide sequence alignment of YRDC promoter regions

```
G11         TTAATAAAGTGGTGGATTAACCTCGCTCGAAAAATGGGGGAAAAAAGGTCGAAAATAAGG
B9          TTAATAAAGTGGTGGATTAACCTCGCTCGAAAAATGGGGGAAAAAAGGTCGAAAATAAGG
A8          TTAATAAAGTGGTGGATTAACCTCGCTCGAAAAATGGGGGAAAAAAGGTCGAAAATAAGG
RCC4222     TTAATAAAGTGGTGGATTAACCTCGCTCGAAAAATGGGGGAAAAAAGGTCGAAAATAAGG
RCC1868     ----------------------------------------------------------GG
B1          -------------------------------------------------------GCGGG
G7          ----------------------------------------------------------GG
                                                                      **

G11         AAAAATATCAACGACGAAC-----------------------------------------
B9          AAAAATATCAACGACGAAC-----------------------------------------
A8          AAAAATATCAACGACGAAC-----------------------------------------
RCC4222     AAAAATATCAACGACGAACGAAAAAGCGAGGCAGAGGAGCTCATATCGTTTCCAATCAGA
RCC1868     AAAAATATCAACGACGAAC-----------------------------------------
B1          AAAAATATCAACGACGAACGAAAAAGCGAGGCAGAGGAGCTCATATCGTTT-CAATCAGA
G7          AAAAATATCAACGACGAAC-----------------------------------------
            *******************
```

```
G11         ------------------------------------------------------------
B9          ------------------------------------------------------------
A8          ------------------------------------------------------------
RCC4222     GAGACCTGAGCGACGACGTGATAAAAAAGTTGACGATGTCCGGACGATGCACGCCGTCGT
RCC1868     ------------------------------------------------------------
B1          GAGACCTGAGCGACGACGTGATAAAAAAGTTGACGATGTCCGGACGATGCACGCCGTCGT
G7          ------------------------------------------------------------


G11         ------------------------------------------------------------
B9          ------------------------------------------------------------
A8          ------------------------------------------------------------
RCC4222     CAGATCCCGCCGCGCAAAGTAGCAGGCGTTTCTTCTTTGTCTTCTTCTTCTTCTTGATCG
RCC1868     ------------------------------------------------------------
B1          CAGATCCCGCCGCGCAAAGTAGCAGGCGTTTCTTCTTTGTCTTCTTCTTCTTCTTGATCG
G7          ------------------------------------------------------------


G11         ----------------------GAAAAACTTTTGCCGTTTCGAGAGGAAATGAAGAAGAT
B9          ----------------------GAAAAACTTTTGCCGTTTCGAGAGGAAATGAAGAAGAT
A8          ----------------------GAAAAACTTTTGCCGTTTCGAGAGGAAATGAAGAAGAT
RCC4222     TCGTCTCTTTTTTTTTCGCGAGGGAAAACTTTTGCCGTTTCGAGAGGAAATGAAGAAGAT
RCC1868     -----------------------GAAAACTTTTGCCGTTTCGAGAGGAAATGAAGAAGAT
B1          TCGTCTCTTTTTTTTTCGCGAGGGAAAACTTTTGCCGTTTCGAGAGGAAATGAAGAAGAT
G7          ----------------------GAAAAACTTTTGCCGTTTCGAGAGGAAATGAAGAAGAT
                                   *************************************

G11         TTTGGGAAACATTTTGCTTCAGTGTGAGGACGCTTTCTCTCTCTCTTTACGCCGAGGACA
B9          TTTGGGAAACATTTTGCTTCAGTGTGAGGACGCTTTCTCTCTCTCTTTACGCCGAGGACA
A8          TTTGGGAAACATTTTGCTTCAGTGTGAGGACGCTTTCTCTCTCTCTTTACGCCGAGGACA
RCC4222     TTTGGGAAACATTTTGCTTCAGTGTGAGGACGCTTTCTCTCTCTCTTTACGCCGAGGACA
RCC1868     TTTGGGAAACATTTTGCTTCAGTGTGAGGACGCTTTCTC-------TTACGCCGAGGACA
B1          TTTGGGAAACATTTTGCTTCAGTGTGAGGACGCTTTCTCTCTCTCTTTACGCCGAGGACA
G7          TTTGGGAAACATTTTGCTTCAGTGTGAGGACGCTTTCTCTCTCTCTTTACGCCGAGGACA
            ***************************************       **************

G11         CTCATCATCATGCGCGCATCATCGACACCGC
B9          CTCATCATCATGCGCGCATCATCGACACCGC
A8          CTCATCATCATGCGCGCATCATCGACACCGC
RCC4222     CTCATCATCATGCGCGCATCATCGACACCG-
RCC1868     CTCATCATCATGCGCGCATCATCGACACCG-
B1          CTCATCATCATGCGCGCATCATCGACACCG-
G7          CTCATCATCATGCGCGCATCATCGACACCG-
```

### B. Marker TOC1 promoter

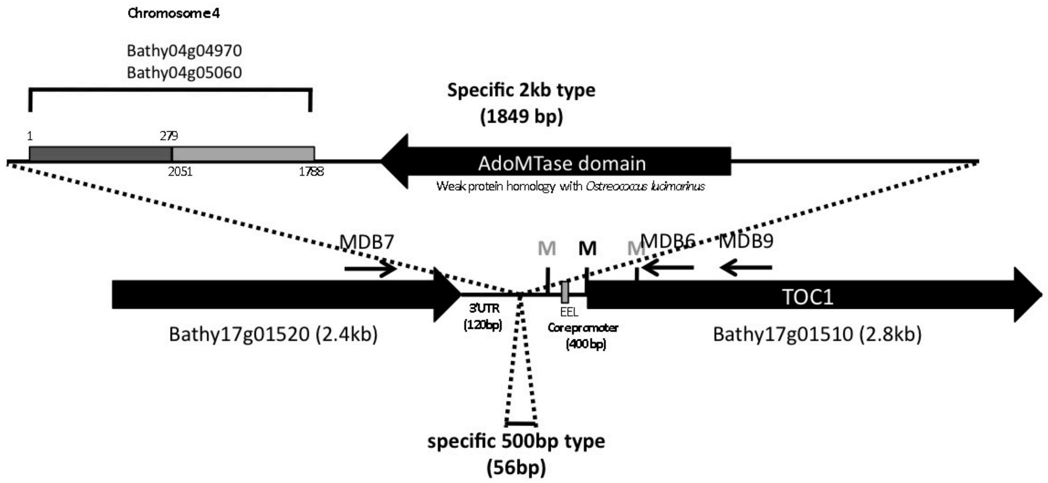


**Organisation of the *TOC1* locus**

Schematic representation of the upstream region of gene Bathy17g01510 encoding an homolog of TOC1 involved in the control of circadian rhythm. RCC1613, RCC1615 and RCC4752 possess a large insertion encoding 2 truncated proteins and one complete AdoMTase and belong to the 2kb insertion type (size amplification with MDB6+MDB7). The other accessions do not have an insertion in comparison to RCC4222 and are grouped into the 500 bp type. The position of the primers used for amplification and sequencing is indicated by arrowheads. Dotted lines indicate insertion or deletion.

Sequencing revealed the presence of an entire potentially functional gene encoding a Methyltransferase-like protein (AdoMTase *METTL24* IPR026913 IPR029063) of 320 aa with 41.7% amino acids identity to the predicted *Ostreococcus lucimarinus* CCE9901 protein (XP_001422352). This gene can be considered as an accessory gene in the genome of *Bathycoccus prasinos*. In addition two truncated genes similar to Bathy04g04970 and Bathy04g05060 encoding similar hypothetical proteins were rearranged at the 5'end of the insertion. The insertion is located upstream the core promoter of *TOC1* containing the essential Evening Element-Like (EEL) *cis* element.

#### Multiple nucleotide sequence alignment of the TOC1 promoter regions

```
RCC1868   ------AAAACAAAAAAACTTTCTTTGACGAGATGAAAAAAGTGCATTTCCCGTGTTCTT      54
RCC5417   -------AAACAAAAAAACTTTCTTTGACGAGATGAAAAAAGTGCATTTCCCGTGTTCTT      53
A8        AAAACAAGAACAAAACTGCTTTTGATGACGCGATGA----------------AAAAAGTG      44
RCC4222   -AAACAAAAACAAAACTGCTTTTGATGACGCGATGA----------------AAAAAGTG      43
B1        AAAACAAAAACAAAACTGCTTTTGATGACGCGATGA----------------AAAAAGTG      44
RCC685    -AAAACAAAAAAAAACTGCTTTTGATGACGCGATGA----------------AAAAAGTG      43
B9        -AAACAAAAAAAAAACTGCTTTTGATGACGC-GGTG----------------AAAAAAGG      42
H11       -AAACAAAAAAAAAACTGCTTTTGATGACGC-GGTG----------------AAAAAAGG      42
RCC1613   AAAACAAAAACAAAACTGCTTT--TGATGAC-GATG----------------AAAAAAGG      41
RCC4752   AAAACAAAAACAAAACTGCTTT--TGATGAC-GATG----------------AAAAAAGG      41
RCC1615   AAAACAAAAAAAAAACTGCTTT--TGATGAC-GATG----------------AAAAAAGG      41
                  ** ****   ****

RCC1868   TTGTGTTGGATGCTTCTGCGTCTCAAATTTCTATTCAAGTGCAAAAGATCTCATAAATAT      114
RCC5417   TTGTGTTGGATGCTTCTGCGTCAAAAATTTCTATTCAAGTGCAAAAGATCTCATAAATAT      113
A8        CATTTCAGGATGCCTCTGCGTCTCAAATTTCTATTCAAGTGCAAAAGATCTCATAAATAT      104
RCC4222   CATTTCAGGATGCCTCTGCGTCTCAAATTTCTATTCAAGTGCAAAAGATCTCATAAATAT      103
B1        CATTTCAGGATGCCTCTGCGTCTCAAATTTCTATTCAAGTGCAAAAGATCTCATAAATAT      104
RCC685    CATTTCAGGATGCCTCTGCGTCTCAAATTTCTATTCAAGTGCAAAAGATCTCATAAATAT      103
B9        CATTTCAGGATGCTTCTGCGTCTCATATTTCTATTCAAGTGCAAAAGATCTCATAAATAT      102
H11       CATTTCAGGATGCTTCTGCGTCTCATATTTCTATTCAAGTGCAAAAGATCTCATAAATAT      102
RCC1613   CATTTCAGGATGCTTCTGCGTCTCAAATTTCTATTCAAGC--AAAAGATCTCATAAATAT      99
RCC4752   CATTTCAGGATGCTTCTGCGTCTCAAATTTCTATTCAAGC--AAAAGATCTCATAAATAT      99
RCC1615   CATTTCAGGATGCTTCTGCGTCTCAAATTTCTATTCAAGTGCAAAAGATCTCATAAATAT      101
             *   ****** ********  * *************   ******************
```

```
RCC1868    CTCAAGAAAAAACCGCTCTTTCGTACGGGAAGGAGGAAACACATTCTC------TAATCA      168
RCC5417    CTCAAGAAAAAAACCGCTCTTCGTCGGGGAAGGAGGAAACACATTATAAATCACACATAA      173
A8         CTCAAGAAAAAACCGCTCTTTACG--GGGAAGGAGGAAACACATTCTCTAATCACA----      158
RCC4222    CTCAAGAAAAAACCGCTCTTTACG--GGGAAGGAGGAAACACATTCTCTAATCACA----      157
B1         CTCAAGAAAAAACCGCTCTTTACG--GGGAAGGAGGAAACACATTCTCTAATCACA----      158
RCC685     CTCAAGAAAAAACCGCTCTTTACG--GGGAAGGAGGAAACACATTCTCTAATCACA----      157
B9         CTCAAGAAAAAACCGCTCTTTCGTACGGGAAGGAGGAAACACATTCTCTAATCACA----      158
H11        CTCAAGAAAAAACCGCTCTTTCGTACGGGAAGGAGGAAACACATTCTCTAATCACA----      158
RCC1613    CTCAAGAAAAAACCGCTCTTTCGTACGGGAAGGAGGAAACACATTCTCTAATCACA----      155
RCC4752    CTCAAGAAAAAACCGCTCTTTCGTACGGGAAGGAGGAAACACATTCTCTAATCACA----      155
RCC1615    CTCAAGAAAAAACCGCTCTTTCGTACGGGAAGGAGGAAACACATTCTCTAATCACA----      157
           ************ *     **     ******************* *

RCC1868    CACAAAGAAGGTCACGAGGAGTGCAGAGAGACAGAGCGCGCACACACACGCGTAAACAAA      228
RCC5417    AAGAAAGAAGGTCACGAGGAGTGCAGAGAGACAGAGCGCGCACACACACGCGTAAACAAA      233
A8         CAAAGAAGGTCACGAG--GAGTGCAGAGAGACAGCGCGCGCACACACACGCGTAAACAAA      216
RCC4222    CAAAGAAGGTCACGAG--GAGTGCAGAGAGACAGCGCGCGCACACACACGCGTAAACAAA      215
B1         CAAAGAAGGTCACGAG--GAGTGCAGAGAGACAGCGCGCGCACACACACGCGTAAACAAA      216
RCC685     CAAAGAAGGTCACGAG--GAGTGCAGAGAGACAGCGCGCGCACACACACGCGTAAACAAA      215
B9         CAAAGAAGGTCACGCGAGGAGTGCAGAGAGACAGCGCGCGCACACACACGCGTAAACAAA      218
H11        CAAAGAAGGTCACGCGAGGAGTGCAGAGAGACAGCGCGCGCACACACACGCGTAAACAAA      218
RCC1613    CAAAGAAGGTCACGAG--GAGTGCAGAGAGACAGAGCGCGCACACACACGCGTAAACAAA      213
RCC4752    CAAAGAAGGTCACGAG--GAGTGCAGAGAGACAGAGCGCGCACACACACGCGTAAACAAA      213
RCC1615    CAAAGAAGGTCACGAG--GAGTGCAGAGAGACAGCGCGCGCACACACACGCGTAAACAAA      215
            * * *      *  *  **************** *************************

RCC1868    GAGAGAGATATACCATGAGGCGATCGACGCGCGCGCACAGATCTTCCTCTGGAAAAGAAG      288
RCC5417    GAGAGAGATATACCATGAGGCGATCGACGCGCGCGCACAGATCTTCCTCTGGAAAAGAAG      293
A8         GAGAGAGATATACCATGAGGCGATCGACGCGCGCGCACAGATCTTCCTCTGGAAAAGAAG      276
RCC4222    GAGAGAGATATACCATGAGGCGATCGACGCGCGCGCACAGATCTTCCTCTGGAAAAGAAG      275
B1         GAGAGAGATATACCATGAGGCGATCGACGCGCGCGCACAGATCTTCCTCTGGAAAAGAAG      276
RCC685     GAGAGAGATATACCATGAGGCGATCGACGCGCGCGCACAGATCTTCCTCTGGAAAAGAAG      275
B9         GAGAGAGATATACCATGAGGCGATCGACGCGCGCGCACAGATCTTCCTCTGGAAAAGAAG      278
H11        GAGAGAGATATACCATGAGGCGATCGACGCGCGCGCACAGATCTTCCTCTGGAAAAGAAG      278
RCC1613    GAGAGAGATATACCATGAGGCGATCGACGCGCGCGCGCAGATCTTCCTCTGGAAAAGAAG      273
RCC4752    GAGAGAGATATACCATGAGGCGATCGACGCGCGCGCGCAGATCTTCCTCTGGAAAAGAAG      273
RCC1615    GAGAGAGATATACCATGAGGCGATCGACGCGCGCGCACAGATCTTCCTCTGGAAAAGAAG      275
           ************************************ ***********************

RCC1868    AGGGAAAAGAAAAGAGGAATCTTGAGTCTCCTTTTCCAATCCGGAAACGGAGTGACGACA      348
RCC5417    AGGGAAAAGAAAAGAGGAATCTTGAGTCTCCTTTTCCAATCCGGAAACGGAGTGACGACA      353
A8         AGGGAAAAGAAAAGAGGAATCTTGAGTCTCCTTTTCCAATCCGGAAACGGAGTGACGACA      336
RCC4222    AGGGAAAAGAAAAGAGGAATCTTGAGTCTCCTTTTCCAATCCGGAAACGGAGTGACGACA      335
B1         AGGGAAAAGAAAAGAGGAATCTTGAGTCTCCTTTTCCAATCCGGAAACGGAGTGACGACA      336
RCC685     AGGGAAAAGAAAAGAGGAATCTTGAGTCTCCTTTTCCAATCCGGAAACGGAGTGACGACA      335
B9         AGGGAAAAGAAAAGAGGAATCTTGAGTCTCCTTTTCCAATCCGGAAACGGAGTGACGACA      338
H11        AGGGAAAAGAAAAGAGGAATCTTGAGTCTCCTTTTCCAATCCGGAAACGGAGTGACGACA      338
RCC1613    AGGGAAAAGAAAAGAGGAATCTTGAGTCTCCTTTTCCAATCCGGAAACGGAGTGACGACA      333
RCC4752    AGGGAAAAGAAAAGAGGAATCTTGAGTCTCCTTTTCCAATCCGGAAACGGAGTGACGACA      333
RCC1615    AGGGAAAAGAAAAGAGGAATCTTGAGTCTCCTTTTCCAATCCGGAAACGGAGTGACGACA      335
           ************************************************************

RCC1868    CGTCGGAGTTTGAGGATGATTTTGTTGAGAAGAAGACTACGACGAAGAAGACGAAGAGGA      408
RCC5417    CGTCGGAGTTTGAGGATGATTTTGTTGAGAAGAAGACTACGACGAAGAAGACGAAGAGGA      413
A8         CGTCGGAGTTTGAGGATGATTTTGTTGAGAAGAAGACTACGACGAATGAGACGAAGAGGA      396
RCC4222    CGTCGGAGTTTGAGGATGATTTTGTTGAGAAGAAGACTACGACGAAGAAGACGAAGAGGA      395
B1         CGTCGGAGTTTGAGGATGATTTTGTTGAGAAGAAGACTACGACGAAGAAGACGAAGAGGA      396
RCC685     CGTCGGAGTTTGAGGATGATTTTGTTGAGAAGAAGACTACGACGAAGAAGACGAAGAGGA      395
B9         CGTCGGAGTTTGAGGATGATTTTGTTGAGAAGAAGACTACGACGATGAAGGCGAAGAGGA      398
H11        CGTCGGAGTTTGAGGATGATTTTGTTGAGAAGAAGACTACGACGAAGAAGGCGAAGAGGA      398
RCC1613    CGTCGGAGTTTGAGGATGATTTTGTTGAGAAGAAGACTACGACGAAGAAGACGAAGAGGA      393
RCC4752    CGTCGGAGTTTGAGGATGATTTTGTTGAGAAGAAGACTACGACGAAGAAGACGAAGAGGA      393
RCC1615    CGTCGGAGTTTGAGGATGATTTTGTTGAGAAGAAGACGAAGAGG---------------A      382
           ************************************* * ** **

RCC1868    CG         410
RCC5417    TG         415
A8         CG         398
RCC4222    TG         397
B1         TG         398
RCC685     TG         397
B9         CG         400
```

```
H11        CG        400
RCC1613    TG        395
RCC4752    TG        395
RCC1615    TG        382
```

### C. Marker Flavodoxin-like protein

The number of repeats in the  flavodoxin-like protein encoded by gene Bathy03g02080 was variable between strains.

#### Multiple amino acid sequence alignment of flavodoxins

```
B1          QAERDAEEARIRAADEEEKARIEAEMKAQEEEEARIKAAEEEERARIEAEMKAKEEEEAR
G7          QAERDAEEARIRAAEEEEKARIEAEMKAQEEEEARIRAAEEEERARIEAEMKAKEEEEAR
C2          QAERDAEEARIRAAEEEEKARIEAEMKAQEEEEARIRAAEEEERARIEAEMKAKEEEEAR
RCC1868     QAERDAEEARIRAAEEEEKARIEAEMKAQEEEEARIRAAEEE------------------
RCC4222     QAERDAEEARIRAAEEEEKARIEAEMKAQEEEEARIRAAEEEERARIEAEMKAKEEEEAR
RCC5417     QAERDVEEARIRAAEEEE------------------------------------------
            *****.********:***

B1          IKAAEEEERARIEAEMKAKEEEEARIKAAEEEERARIEAEMKAQEEEEARIKAAEEEERA
G7          IKAAEEEERARIEAEMKAQEEEEARIKAAEEEERARIEAEMKAQEEEEARIKAAEEEERA
C2          IRAAEEEERARIEAEMKAQEEEEARIKAAEEEERARIEAEMKAQEEEEARIKAAEEEERA
RCC1868     -------ERARIEAEMKAQEEEEARIKAAEEEERARIEAEMKAKEEEEARIKAAEEEERA
RCC4222     IKAAEEEERARIEAEMKAQEEEEARIKAAEEEERARIEAEMKAKEEEEARIKAAEEEERA
RCC5417     ---------------------------------RARIEAEMKAKEEEEARIRAAEEEERA
                                                        ******:********

B1          RIEAEMKAQEEEEARIKAAEEEERARIEAEMKAQEEEEARIKAAEEEERARIEAEMKAKE
G7          RIEAEMKAKEEEEARIKAAEEEERARIEAEMKAQEEEEARIKAAEEEERARIEAEMKAQE
C2          RIEAEMKAKEEEEARIKAAEEEERARIEAEMKAQEEEEARIKAAEEEERARIEAEMKAQE
RCC1868     RIEAEMKAQEEEEARIKAAEEEERARIEAEMKAKEEEEARIKAAEEEERARIEAEMKAKE
RCC4222     RIEAEMKAQEEEEARIKAAEEEERARIEAEMKAKEEEEARIKAAEEEERARIEAEMKAKE
RCC5417     RIEAEMKAQEEEEARIRAAEEEERARIEAEMKAQEEEEARIRAAEEEERARIEAEMKAKE
            ********:*******:****************:*******:****************:*

B1          EEEARIKAAEEEERARIEAEMKAKEEEEARIKAAEEEERARIEAEMKAKEEEEARIKAAE
G7          EEEARIKAAEEEERARIEAEMKAKEEEEARMKAAEEEERARIEAEMKAKEEEESRIKAAE
C2          EEEARIKAAEEEERARIEAEMKAKEEEEARMKAAEEEERARIEAEMKAKEEEEARIKAAE
RCC1868     EEEARIKAAEEEERARIEAEMKAKEEEDA-------------------------RIKAEE
RCC4222     EEEARIKAAEEEERARIEAEMKAKEEEDA-------------------------RIKAEE
RCC5417     EEEARIRAAEEEEKAR---------------------------------------INAEE
            ******:******:**                                       *:* *

B1          EEERARIEAEMKAKEEEEARIKAAEEEERARIEAEMKAKEEEEARIKAAEEEERARIEAE
G7          EEERARIEAEMKAKEEEESRIKAAEEVVRASIVSCMKAK---------------------
C2          EEERARIEAEMKAKEEEEARIKAAEEEERARIEAEMKAKEEEEARMKAAEEEERARIEAE
RCC1868     EE-KARIEAEQKAIAAAEANRAA-------------------------------------
RCC4222     EE-EARIEAEQKAIAAAEANRAA-------------------------------------
RCC5417     EE-KARIEAEQKAIAAAEANRAA-------------------------------------
            ** .****** **    *:.  *

B1          MKAKEEEEARIKAAEEEERARIEAEMKAKEEEEARIKAAEEEERARIEAEMKAKEEEEAR
G7          ------------------------------------------------------------
C2          MKAKEEEEARIKAAEEEERARIEAEMKAKEEEESI-------------------------
RCC1868     ------------------------------------------------------------
RCC4222     ------------------------------------------------------------
RCC5417     ------------------------------------------------------------


B1          IKAAEEEERARIEAEMKAKEEEESSMKAAELVV
G7          ---------------------------------
C2          ---------------------------------
RCC1868     ---------------------------------
```

```
RCC4222      ---------------------------------
RCC5417      ---------------------------------
```

### D. Marker Zinc finger

The number of repeats in the zinc finger protein (C2H2 Family) encoded by Bathy15g02300 was variable between strains

#### Multiple amino acid sequence alignment of Zinc finger proteins

```
RCC5417      MSNRRKNPHPKRAENVKTVLKGEKEER-PLLLCLRIPDEEDD----DDDVDFDEVVEKQN     55
RCC685       MSSRRKNPHPKRAENARTVLKGEKEER-PVLLCVRIPDDDDDEDDDEEDVAFDEVVEQQN     59
RCC1613      MSSRRKNPHPKRAENARTVLKGEKEER-PVLLCVRIPDDDDDEDDDEEDVAFDEVVEQQN     59
G11          MSNRRKNPHPKRAENVKTVLKGEKEER-PVLLCVRIPDDDD-EDDDEEDVAFDEVVEQQN     58
RCC1105      MSSRRKNPHPKRAENARTVLKGEKEER-PVLLCVRIPDDDD-EDDDEEDVAFDEVVEQQN     58
G7           MSNRRKNPHPKRAENVKTVLKGEKEER-PVLLCVRIPDDDD-EDDDEEDVAFDEVVEQQN     58
B9           MSNRRKNPHPKRAENVKTVLKGEKEER-PVLLCVRIPDDDD-EDDDEEDVAFDEVVEQQN     58
RCC1868      MSSRRKNPHPKRAENARTVLKGEKEERPPVLLCVRIPDDDDDEDDDEEDVAFDEVVEQQN     60
RCC4752      MSSRRKNPHPKRAENARTVLKGEKEER-PVLLCLRIPDDDD-EDDDEEDVAFDEVVEQQN     58
A8           MSSRRKNPHPKRAENARTVLKGEKEER-PVLLCVRIPDDDD-EDDDEEDVAFDEVVEQQN     58
             **.************.:********** *:***:****::*     ::** ******:**

RCC5417      EKMRKEKTATKRGGKRKRGQQHECDVCEKMFGYASQLAIHMRIHTNEKPYECDVCEKRFR     115
RCC685       EKLRKEKTATKRGGKRKRGQQHECDLCEKVFDRPSSLARHMRIHTNERPYECDVCEMRFR     119
RCC1613      EKLRKEKTATKRGGKRKRGQQHECDLCEKVFDRPSSLAIHMRIHTNEKPYECDVCEMRFR     119
G11          EKLRKEKTATKRGGKRKRGQQHECDVCEKVFDRPSSLAMHMRIHTNEKPYECDVRD-SFR     117
RCC1105      EKLRKEKTATKRGGKRKRGQQHECDVCEKVFDRPSSLAIHMRIHTNEKPYECDVCEMRFR     118
G7           EKLRKEKTATKRGGKRKRGQQHECDVCEKVFDRPSSLAMHMRIHTNEKPYECDVCEMRFR     118
B9           EKLRKEKTATKRGGKRKRGQQHECDVCEKVFDRPSSLAMHMRIHTNEKPYECDVCEMRFR     118
RCC1868      EKLRKEKTATKRGGKRKRGQQHECDVCEKVFDRPSSLAIHKRIHTNEKPYECDVCEKSFS     120
RCC4752      EKLRKEKTATKRGGKRKRGQQHECDVCEKVFDRPSSLAIHKRIHTNEKPYECDVCEKSFS     118
A8           EKLRKEKTATKRGGKRKRGRQHECDLCEKVFDRPSSLAIHMRIHTNEKPYECDVCEKSFS     118
             **:****************:*****:***:*.  *.** * ******:****** :  *

RCC5417      ESGDLKKHKRIHTKEKPYECDVCEMRFRTSDVLKTHMRIHTNEKRYECAVCEKRYRHSSA     175
RCC685       HSSTLRVHKRIHTNEKPYECDICDKAFRESGKLKEHMRIHTNEKPNECDVCEKRFTQ---     176
RCC1613      HSSTLRVHKRIHTNEKPYECDICDKAFRESGKLKEHMRIHTNEKPYECDNSEKRFSR---     176
G11          HSSTLRVHKRIHTNEKPYECDICDKAFRESGKLKEHMRIHTNEKPYECDVCEKRFSR---     174
RCC1105      HSSTLRVHERIHTNEKPYECDICDKAFRESGKLKEHMRIHTNEKPYECDVCEKRFSR---     175
G7           HSSTLRVHKRIHTNEKPYECDICDKAFRESGKLKEHMRIHTNEKPYECDVCEKRFSR---     175
B9           HSSTLRVHKRIHTNEKPYECDICDKAFRESGKLKEHMRIHTNEKPYECDVCEKRFSR---     175
RCC1868      TAGNLKVHMRIHTNEKPYECDVCEKRFRESSTLQNHMRIHTNEKPYECDVCEKRFSR---     177
RCC4752      TAGNLKVHMSIHTNEKPYECDVCEKRFRESGTLQNHMRIHTNEKPYECDVCEKRFSR---     175
A8           TAGNLKVHMRIHTNEKPYECDVCEKRFRESATLQNHMRIHTNEKPYECDVCEKRFSR---     175
             :. *: *  ***:*******:*:  ** *  *: *********  **  .***: :

RCC5417      LKSHMRIHTNEMFECDVCQKRFTLASNLKRHMLIHTNEKPYECDVCEMRFTQPDNLKRHK     235
RCC685       ------------------------TANLKKHMLIHTNEKPYECDVCEMRFRQSQHLKAHK     212
RCC1613      ------------------------SGTLQSHMRIHTNEKPYECDVCEKRFRTSSQLKVHV     212
G11          ------------------------SGTLQSHMRIHTNEKPYECDVCEKRFRTSSQLKVHV     210
RCC1105      ------------------------SGTLQSHMRIHTNEKPYECDVCEKRFRTSSQLKVHV     211
G7           ------------------------SGTLQSHMRIHTNEKPYECDVCEKRFRTSSQLKVHV     211
B9           ------------------------SGTLQSHMRIHTNEKPYECDVCEKRFRTSSQLKVHV     211
RCC1868      ------------------------SGTLQSHMRIHTNEKPYECDVCEKRFRTSGQLKVHV     213
RCC4752      ------------------------SGTLQSHMRIHTNEKPYECDVCEKRFRTSSQLKEHM     211
A8           ------------------------SGTLQSHMRTHR------------------------     187
                                    :..*: **  *

RCC5417      RIHTKEKPYECDVCEKRFRESSDLKKHMRTHTNEKPYECDVCDKAFRNSGQLKVHMRIHT     295
RCC685       RIHTNEKPYECDVCEKRFSESGALKSHVRIHTNEKPYECDVCEKCFRHSSALRVHKRTHR     272
RCC1613      RIHTNEKAYECDVCEKRFTQSCNLKTHMRTHR----------------------------     244
G11          RIHTNEKAYECDVCEKRFTQSCNLKTHMRTHR----------------------------     242
RCC1105      RIHTNEKAYECDVCEKRFTQSCNLKTHMRTHR----------------------------     243
G7           RIHTNEKAYECDVCEKRFTQSCNLKTHMRTHR----------------------------     243
B9           RIHTNEKAYECDVCEKRFTQSCNLKTHMRTHR----------------------------     243
RCC1868      RIHTNEKAYECDVCEKRFTQSGSLKYHMRTQH----------------------------     245
RCC4752      RIHTNEKAYECDVCEKRFTQSCNLKTHMRTHR----------------------------     243
```

```
A8          ------------------------------------------------------------     187

RCC5417     NEKPYECDVCVKRFTQSSNLKRHVFCCH        323
RCC685      ----------------------------        272
RCC1613     ----------------------------        244
G11         ----------------------------        242
RCC1105     ----------------------------        243
G7          ----------------------------        243
B9          ----------------------------        243
RCC1868     ----------------------------        245
RCC4752     ----------------------------        243
A8          ----------------------------        187
```

### E. Marker TIM a

The Tima protein is encoded by Bathy14g03100 on Chromosome 14.

#### Multiple amino acid sequence alignment of Tima proteins

```
RCC4752         MFSPFFEVKLIAFLTLPTEKSATNVQEQNRAHLNDLNALIENENIFAIVLGLILEPLKQL 60
RCC685          MFSPFFEVKLIAFLTLPTEKSATNVQEQNRAHLNDLNALIENENIFAIVLGLILEPLKQL 60
RCC4222         MFSPFFEVKLIAFLTLPTEKSATNVQEQNRAHLNDLNALIENENIFAIVLGLILEPLKQL 60
RCC1613         MFSPFFEVKLIAFLTLPTEKSATNVQEQNRAHLNDLNALIENENIFAIVLGLILEPLKQL 60
RCC5417         MFSPFFEVKLIAFLTLPTEKSATNVQEQNRAHLNDLNALIENENIFAIVLGLILEPLKQL 60
                                 *****************:********:*********:***:**

RCC4752         ESSGGANFK-TNESKSLQLILTFYRNLLIIAEEENNTRVKKILSSCLFKYNFLDVLLVLV 119
RCC685          ESSGGANFK-TNESKSLQLILTFYRNLLIIAEEENNTRVKKILSSCLFKYNFLDVLLVLV 119
RCC4222         ESSGGANFK-TNESKSLQLILTFYRNLFIIAEEENNTRVKKILSSCLFKYNFLDVLLVLV 119
RCC1613         ESSGGANFK-TNESKSLQLILTFYRNLLIIAEEENNTRVKKILSSCLFKYNFLDVLLVLV 119
RCC5417         ESSGGANFK-TNESKSLQLILTFYRNLLIIAEEENNTRVKKILSSCLFKYNFLDVLLVLV 119
                ** ***::  ..*******:*******:************:******::*.*********

RCC4752         QNHKKILGHEDTALAVEILQLLFRGIPPDKLAEELSNAQNKDQDIFTHDSLKFPYQTKKH 179
RCC685          QNHKKILGHEDTALAVEILQLLFRGIPPDKLAEELSNAQNKDQDIFTHDSLKFPYQTKKH 179
RCC4222         QNHKKILGHEDTALAVEILQLLFRGIPPDKLAEELSNAQNKDQDIFTHDSLKFPYQTKKH 179
RCC1613         QNHKKILGHEDTALAVEILQLLFRGIPPDKLAEELSNAQNKDQDIFTHDSLKFPYQTKKH 179
RCC5417         QNHKKILGHEDTALAVEILQLLFRGIPPDKLAEELSNAQNKDQDIFTHDSLKFPYQTKKH 179
                *******.*** **::*********** : :  .* .*:: .:*  .  .**:  : * :

RCC4752         IPRFNGKFEPKSVRRVV-EMSGKIKKSHIHPVLCPEKSYFIDKSIPQFWVALLDSTYFGA 238
RCC685          IPRFNGKFEPKSVRRVV-EMSGKIKKSHIHPVLCPEKSYFIDKSIPQFWVALLDSTYFGA 238
RCC4222         IPRFNGKFEPKSVRRVV-EMSGKIKKSHIHPVLCPEKSYFIDKSIPQFWVALLDSTYFGA 238
RCC1613         IPRFNGKFEPKSVRRVV-EMSGKIKKSHIHPVLCPEKSYFIDKSIPQFWVALLDSTYFGA 238
RCC5417         IPRFNGKFEPKSVRRVV-EMSGKIKKSHIHPVLCPEKSYFIDKSIPQFWVALLDSTYFGA 238
                  **:****  *: :    *.*.:...: : .   *.**::*..   **  **:**:**:

RCC4752         FIFHAWRDIVRTSGDVEQRATEWQTRSYNLLRFSTFGLRILSKFF-HKNALKDKISIKCV 297
RCC685          FIFHAWRDIVRTSGDVEQRATEWQTRSYNLLRFSTFGLRILSKFF-HKNALKDKISIKCV 297
RCC4222         FIFHAWRDIVRTSGDVEQRATEWQTRSYNLLRFSTFGLRILSKFF-HKNALKDKISIKCV 297
RCC1613         FIFHAWRDIVRTSGDVEQRATEWQTRSYNLLRFSTFGLRILSKFF-HKNALKDKISIKCV 297
RCC5417         FIFHAWRDIVRTSGDVEQRATEWQTRSYNLLRFSTFGLRILSKFF-HKNALKDKISIKCV 297
                **:************::*********.****:*** **:****:: :..  * : **:.*

RCC4752         ADIFDSSFVHWLRMEWIALENKKDYYGANIVCCMLVEITSIVYNISVHGTSAEKDASKFL 357
RCC685          ADIFDSSFVHWLRMEWIALENKKDYYGANIVCCMLVEITSIVYNISVHGTSAEKDASKFL 357
RCC4222         ADIFDSSFVHWLRMEWIALENKKDYYGANIVCCMLVEITSIVYNISVHGTSAEKDASKFL 357
RCC1613         ADIFDSSFVHWLRMEWIALENKKDYYGANIVCCMLVEITSIVYNISVHGTSAEKDASKFL 357
RCC5417         ADIFDSSFVHWLRMEWIALENKKDYYGANIVCCMLVEITSIVYNISVHGTSAEKDASKFL 357
                ****:***:***********:*******.:**  :*** **: ::** **..***.*:**

RCC4752         IRELYFKSKTESLLEQVCISLKSFKSKYPSAYLFSLINLLSSSLRCCKTHQSQ--YEGIR 415
RCC685          IRELYFKSKTESLLEQVCISLKSFKSKYPSAYLFSLINLLSSSLTITKTHQSQ--YEGIR 415
RCC4222         IRELYFKSKTESLLEQVCISLKSFKSKYPSAYLFSLINLLSSSLRCCKTHQSQ--YEGIR 415
RCC1613         IRELYFKSKTESLLEQVCISLKSFKSKYPSAYLFSLINLLSSSLRCCKTHQSQ--YEGIR 415
RCC5417         IRELYFKSKTESLLEQVCISLKSFKSKYPSAYLFSLINLLSSSLRCCKTHQSQ--YEGIR 415
                ****:**:*:******.* ***. ..*:**:**.*** :*.**:.* * :*.    : :*

RCC4752         KYGNIHDNSIIYYHMRLLKGRNKNESLLGSVYIYLRSLVSANCVERLFSFRHLDCLHDYC 475
```

```
RCC685          KYGNIHDNSIIYYHMRLLKGRNKNESLLGSVYIYLQSLVSANCVERLFSFRHLDCLHDYC 475
RCC4222         KYGNIHDNSIIYYHMRLLKGRNKNESLLGSVYIYLQSLVSANCVERLFSFRHLDCLHDYC 475
RCC1613         KYGNIHDNSIIYYHMRLLKGRNKNESLLGSVYIYLQSLVSANCVERLFSFRHLDCLHDYC 475
RCC5417         KYGNIHDNSIIYYHMRLLKGRNKNESLLGSVYIYLQSLVSANCVERLFSFRHLDCLHDYC 475
                .  ** *:*** :*:::**..*  *::*.**  **:.**.*. **:**:**** ***.:

RCC4752         VMSRREPEHISEYALLLQQECHFVLNCFHLAVTRPAEHLNEKKRLFLEVLFS         527
RCC685          VMSRREPEHISEYALLLQQECHFVLNCFHLAVTRPAEHLNEKKRLFLEVLFS         527
RCC4222         VMSRREPEHISEYALLLQQECHFVLNCFHLAVTRPAEHLNEKKRLFLEVLFS         527
RCC1613         VMSRREPEHISEYALLLQQECHFVLNCFHLAVTRPAEHLNEKKRLFLEVLFS         527
RCC5417         VMSRREPEHISEYALLLQQECHFVLNCFHLAVTRPAEHLNEKKRLFLEVLFS         527
                * *: :*:. :  :  *:: *:******  ::.:  :: : ::::**:: .*
```
